# Discovery of an isoflavone oxidative catabolic pathway in legume root microbiota

**DOI:** 10.1101/2023.08.07.552369

**Authors:** Noritaka Aoki, Tomohisa Shimasaki, Wataru Yazaki, Tomoaki Sato, Masaru Nakayasu, Akinori Ando, Shigenobu Kishino, Jun Ogawa, Sachiko Masuda, Arisa Shibata, Ken Shirasu, Kazufumi Yazaki, Akifumi Sugiyama

**Affiliations:** Research Institute for Sustainable Humanosphere, Kyoto University, Uji, Japan; Riken BioResource Research Center, Tsukuba, Ibaraki, Japan; Division of Applied Life Sciences, Graduate School of Agriculture, Kyoto University, Kyoto, Japan; Plant Immunity Research Group, RIKEN Center for Sustainable Resource Science, Yokohama, Kanagawa, Japan

## Abstract

Isoflavones are major specialized metabolites found in legume plants, where they contribute to environmental adaptation. Isoflavones also play a role human health as promising therapeutic agents. This metabolite group is involved in interactions with soil microorganisms as initiation signals in rhizobial symbiosis and as modulators of the legume root microbiota. We previously reported that isoflavones enrich the Comamonadaceae, a predominant bacterial family in soybean roots, and that microorganisms in legume rhizosphere soil degrade isoflavones. However, the isoflavone catabolism pathway that underly the isoflavone-mediated legume–microbiota interactions have not yet been clarified. Here, we isolated *Variovorax* sp. strain V35, member of the Comamonadaceae that harbors isoflavone-degrading activity, from soybean roots and discovered a gene cluster responsible for isoflavone degradation named *ifc*. Strain V35 metabolizes isoflavones in a completely distinct oxidative manner from the reductive isoflavone metabolism pathway elucidated in the gut microbiota, in which resulting products enter the tricarboxylic acid cycle. The characterization of *ifc* mutants and heterologously expressed IFC enzymes revealed that isoflavones are catabolized via A-ring cleaving fission, which starts with hydroxylation at the 8-position of the A-ring. We further demonstrated that *ifc* genes are frequently found in bacterial strains isolated from legume plants, including mutualistic rhizobia, and contribute to detoxification of the antibacterial activity of isoflavones. Taken together, our findings reveal an oxidative catabolism pathway of isoflavone in the soybean root microbiota, providing molecular insights into isoflavone-mediated legume–microbiota interactions.

**Significance:** Isoflavones play pivotal roles in plant-environment interactions and in the maintenance and improvement of human health. Bacterial metabolism is a fundamental component of isoflavone-mediated interkingdom interactions. In the human gut, intestinal bacteria convert isoflavones into equol, a highly bioactive compound. However, the fate of isoflavones in the legume rhizosphere has not been elucidated, despite them being the key signaling molecules for nodule symbiosis and modulation of the legume root microbiota. Here, we discovered a novel isoflavone catabolism pathway in the soybean root microbiota and demonstrated the strong association between bacterial catabolic abilities and their interactions with host plants. Collectively, our findings provide new insights into bacterial isoflavone metabolism and a molecular understanding of legume-microbiota interactions.

## Introduction

Plants produce a vast array of low molecular weight organic compounds, known as plant specialized metabolites (PSMs). These metabolites are not directly involved in plant development and reproduction but largely contribute to plant– environment interactions and adaptation (1, 2). PSMs also exhibit a variety of biological activities that are beneficial to human health (3–5). Over the past decade, it has been demonstrated that PSMs are crucial for the interactions between eukaryotic hosts and their associated microbial communities, particularly plant microbiota (6–9).

The bacterial metabolic capacity of PSMs plays a key role in PSM– mediated host-microbiota interactions. Root microbiota members have the ability to degrade and/or utilize host–specific PSMs, conferring competitive advantages on their root colonization and contributing to the assemblage of species-specific root microbiota (10, 11). In the human gut, the gut microbiota detoxify harmful dietary PSMs or convert them into higher bioactive compounds. *Bacteroides xylanisolvens*, for example, degrades nicotine, a toxic alkaloid produced by tobacco, alleviating non-alcoholic fatty liver disease triggered by nicotine accumulation (12). It is also reported that the human gut microbiota activate glucosinolates, a predominant PSM group in the family Brassicaceae, by converting them to isothiocyanate, which exhibits anticancer activity (13). Therefore, discovering the bacterial catabolic pathways of PSMs is important not only for elucidating the mechanisms of host–microbe interactions but also for the industrial and therapeutical application of PSMs.

Isoflavones, a subgroup of flavonoids predominantly found in *Fabaceae* (14), play roles in the establishment of nodule symbiosis and the assembly of root microbiota (15). Daidzein, a major isoflavone in soybean (*Glycine max*), is secreted into the rhizosphere in response to nitrogen deficiency (16), thereby inducing the expression of *nod* genes of the mutualistic rhizobia, *Bradyrhizobium japonicum* (17, 18), and modulating the soybean root microbiota (19, 20). In addition, its physiological importance in soybean, daidzein is also considered a potential therapeutic agent, since equol, a daidzein-derived metabolite produced by intestinal bacteria, exhibits great estrogenic and antioxidant activity (21, 22), which is already availabe in the market.

In the human gut, daidzein is metabolized by intestinal bacteria and converted to equol or *O*-desmethylangolensin (*O*-DMA) (23). In this pathway, daidzein is metabolized to dihydrodaidzein by a racemase and a reductase. The resulting dihydrodaidzein is further metabolized to equol by a series of reductase reactions or converted to *O*-DMA via the fission of the heterocyclic C-ring. Genes involved in human gut daidzein catabolism have been identified from several intestinal bacteria, including *Bifidobacterium pseudolongum*, *Clostridium* sp., and *Slackia isoflavoniconvertens* (24–27), and differences in the composition of these bacterial species are thought to be responsible for the metabolic variation of intestinal daidzein (28).

In contrast, no genes involved in isoflavone catabolism have been identified in soil microorganisms, although certain rhizobia species degrade isoflavones in a manner distinct from human gut microorganisms (29), suggesting a novel isoflavone metabolic pathway in plant microbiota. We previously demonstrated that microorganisms in the soybean rhizosphere degrade daidzein (30) and that the addition of daidzein to soil led to the enrichment of Comamonadaceae, a predominant bacterial family in soybean roots (20). Comamonadaceae harbors genomic features that enable the catabolism of an array of aromatic compounds (31), implying that these bacterial species are potentially responsible for isoflavone degradation in soybean roots.

In this study, we isolated the isoflavone-degrading soil bacteria, *Variovorax* sp. belonging to the Comamonadaceae, from soybean roots and identified the gene cluster responsible for isoflavone catabolism, which we named “the isoflavone catabolism gene” (*ifc*). By integrating comparative genome analysis and an *in vitro* fitness assay, we demonstrated that the *ifc* gene cluster underlies the interactions between legumes and their root microbiota, including symbiotic mutualists.

## Results

### Isolation of soybean root-associated bacteria involved in isoflavone catabolism

We previously reported enrichment of the family Comamonadaceae in soil treated with daidzein (20) and isolated two Comamonadaceae bacterial genera, *Variovorax* and *Acidovorax*, from soybean roots (Fig. 1a) (32). Among these isolates, seven strains degraded daidzein (Fig. 1a and S1). Phylogenetic analysis based on the 16S rRNA gene sequence revealed that *Variovorax* isolates were closely related to *V. paradoxus* (Fig. 1a). Daidzein-degrading strains were found regardless of their phylogenic distance. Due to its high mutagenesis efficiency, we selected strain V35 for further analysis.

**Fig. 1.**
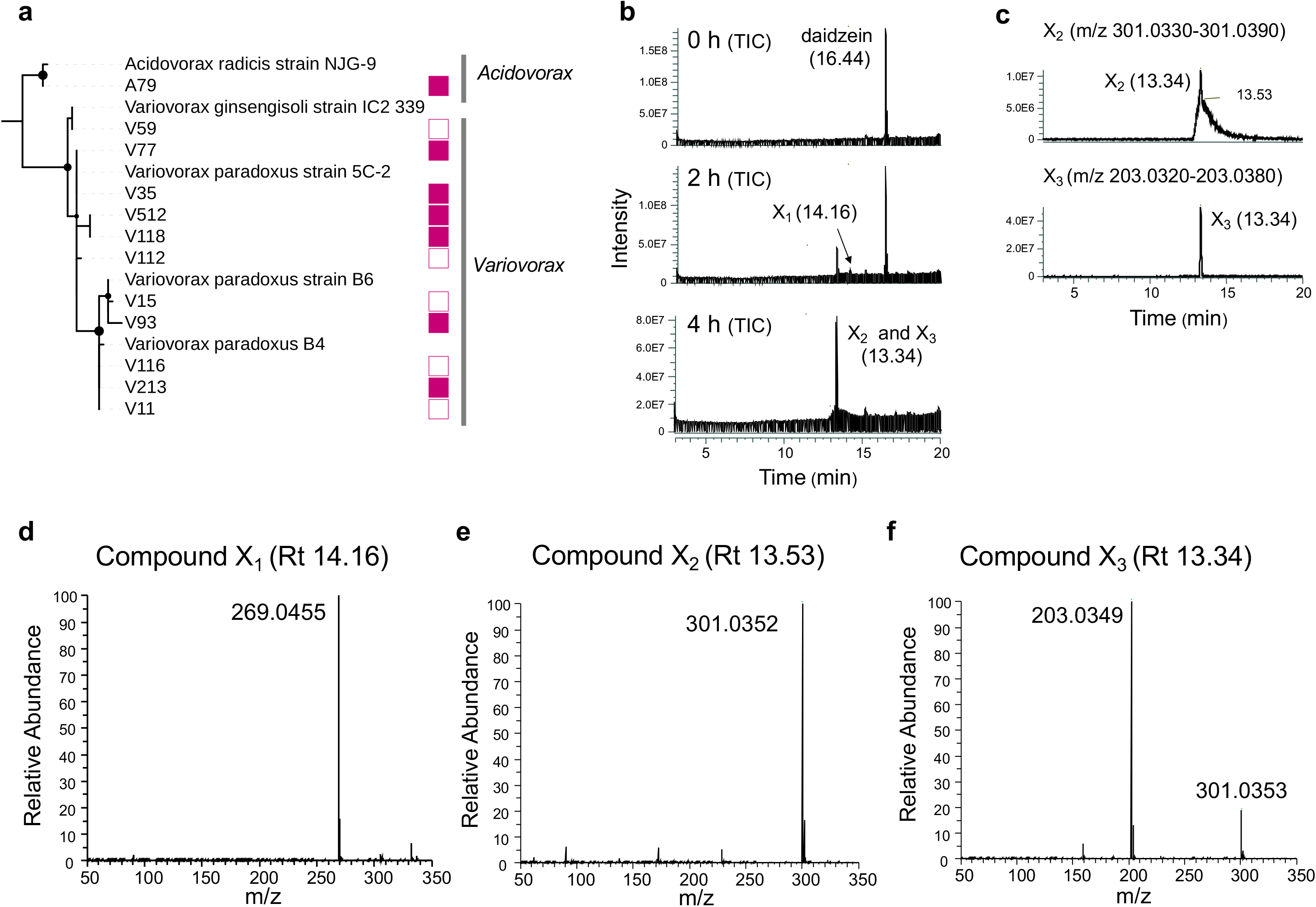
Phylogenetic analysis of soybean root-associated bacteria belonging to family Comamonadaceae and daidzein catabolism intermediates in *Variovorax* sp. strain V35. (a) The phylogenic tree was constructed using the maximum likelihood estimation (MLE) method. Bootstrap values (1,000 replicates) above 0.8 are shown in nodes. The daidzein degradation ability of each isolate is represented by filled and empty squars, respectively. **(b)** LC-MS analysis of the culture supernatant at 0, 2, and 4 h from the V35 culture supernatant grown in daidzein supplemented MS medium. Total ion chromatograms obtained in negative ionization mode with full-scan range of *m/z* 50–350 are shown. **(c)** Selected ion chromatograms of the 4-h culture supernatant obtained in negative ionization mode. Compounds X_2_ and X_3_ were detected with an *m/z* range of 301.0330–301.0390 and m/z 203.0320–203.0380, respectively. **(d)** The mass spectra of compound X_1_-X_3_. Essentially identical results were obtained in two independent experiments.

To identify the daidzein degradation intermediates, we analyzed time-course daidzein degradation of strain V35 by high-resolution mass spectrometer. This strain completely degraded daidzein 4 h after inoculation and produced three reaction product candidates (compounds X_1_–X_3_, Fig. 1b). Total ion current chromatogram analysis in the range of mass-to-charge ratio (*m/z*) of 50–350 detected two peaks in the supernatant at the retention times (Rt) of 13.34 and 14.16. The compound at Rt 14.16 (compound X_1_) showed an ion at *m/z* 269.0455 (Fig. 1d and 2 Sa), consistent with the calculated mass of C_15_H_10_O_5_ (269.0450), which can be formed by mono-hydroxylation of daidzein, suggesting that this compound may be the first reaction product in the daidzein catabolism pathway of this strain. The peak observed at Rt 13.34 consisted of a sharp peak and a broad peak (compounds X_2_ and X_3_, Fig. 1c), with mass fragment ions at *m/z* 203.0349 and 301.0352, respectively (Fig. 1e and f, and S2b and c). Compound X_3_ with an *m/z* of 203.0349 had a lower molecular weight than the other two intermediates, suggesting that this compound is a late intermediate.

**Fig. 2.**
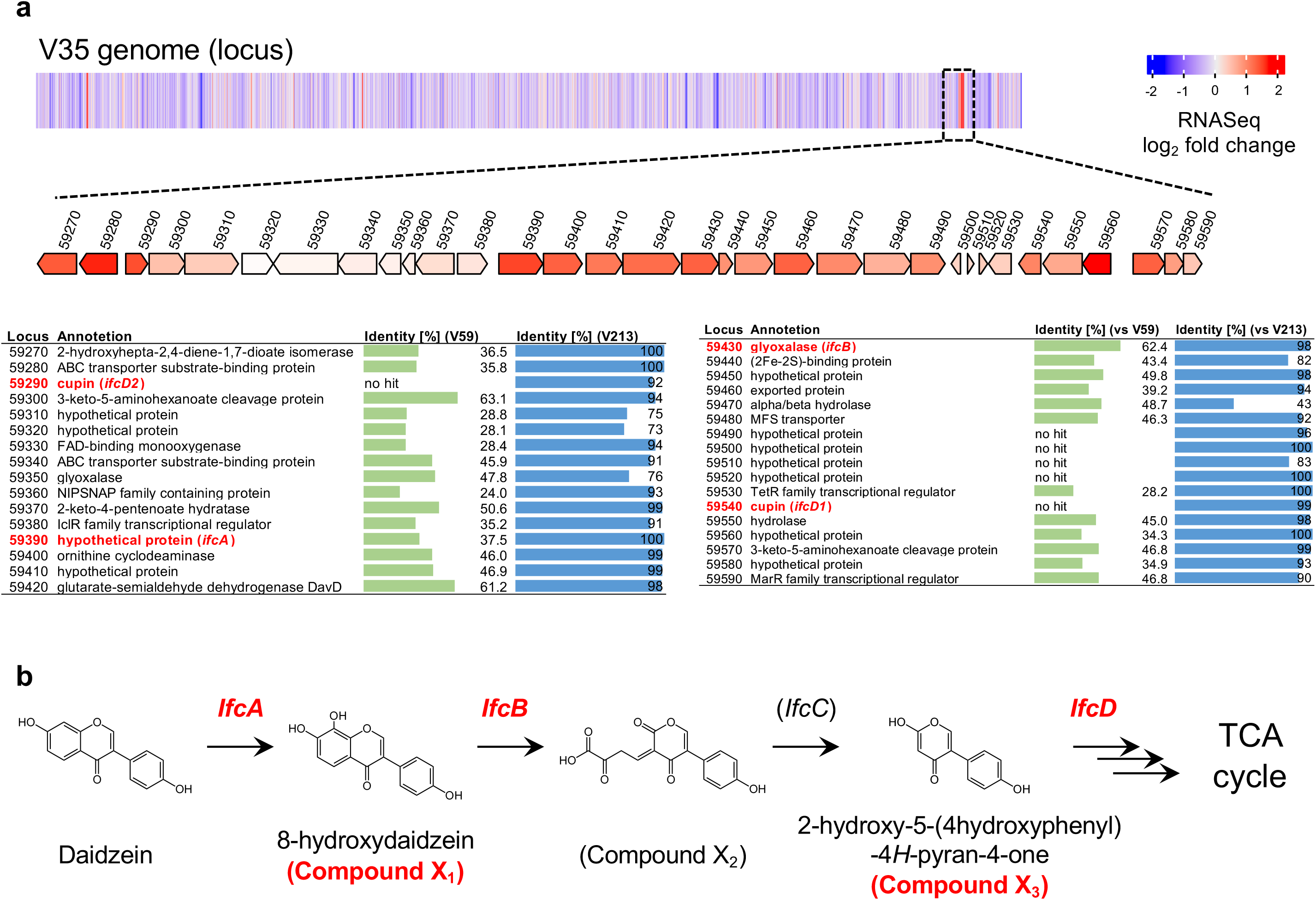
Screening of isoflavone catabolism gene cluster and proposed isoflavone catabolism pathway of genus *Variovorax*. **(a)** A heatmap of the RNA-seq experiment was denoted on the linear chromosome map of strain V35, colored by the log_2_-transformed-fold change in gene expression during the growth in presence of daidzein versus absence of daidzein. The detected hotspot region is displayed on the gene map colored by the log_2_-transformed-fold change (shown in **a**). The functional assignments annotated are shown in the table below the gene map. Per cent identity of each *ifc* gene to the genes from the isoflavone-degrading strain (V213) and the non-degrading strain (V59) are represented in the bar chart. Characterized *ifc* genes are indicated in red. **(b)** Isoflavones were oxidatively catabolized by *ifc* genes and eventually entered the citrate cycle. *Ifc* genes and isoflavone catabolism intermediates identified in this study are represented in red.

### Identification of the isoflavone catabolism gene cluster

To identify isoflavone catabolism genes, we sequenced the genomes of *Variovorax* isolates using the PacBio Sequel II platform and obtained nine complete genome sequences, ranging from one to three closed contigs per strain, and one near complete genome (Dataset S1A). In addition, we carried out transcriptome analysis of V35 grown in the presence or absence of daidzein, because the daidzein degradation activity of V35 was induced by pre cultured in the presence of daidzein (Fig. S3). By integrating comparative transcriptome and genome analysis, we screened for the candidate genes that were 1) upregulated in the presence of daidzein, 2) highly conserved in the daidzein-degrading strains, V35 and V213, and 3) less conserved in the daidzein non-degrading strain, V59. This analysis identified one hotspot (from gene locus 59270 to locus 59590) that was composed of catabolism-related genes such as oxidoreductases, hydrolases, transcriptional regulators, and transporters (Fig. 2a). All genes in the hotspot were upregulated in the presence of daidzein, and 32 out of 33 genes showed higher sequence identity to V213 than to V59. We then hypothesized that this hotspot was a putative gene cluster for isoflavone catabolism, thus designated as *isoflavone catabolism* (*ifc*).

Within the *ifc* cluster, we selected five candidate genes that were potentially involved in the early stage of isoflavone degradation (Fig. 2a). Locus 59390 and locus 59430 were predicted to encode a putative FAD-binding monooxygenase and a putative glyoxalase, respectively. These proteins shared amino acid sequence similarities to *fdeE* (36.3%) and *fdeC* (65.8%), respectively, both of which are involved in the hydroxylation and dioxygenation of the A-ring of naringenin in *Herbaspirillum seropedicae* SmR1 (Fig. S4) (33, 34). Locus 59470 encodes alpha/beta hydrolase and showed sequence similarity with 2,6-dihydropseudooxynicotine hydrolase from *Arthrobacter nicotinovorans* (Q93NG6), which cleaves a C–C bond in 2,6-dihydroxypseudooxynicotine (35). Both locus 59540 and locus 59290 encode proteins belonging to the cupin superfamily, which is characterized by a conserved barrel domain and consists of functionally diverse proteins such as isomerases, oxidoreductases, and seed storage proteins (36). In addition, locus 59540 and locus 59290 shared 65% sequence identity, suggesting that these enzymes are homologous proteins.

### Identification of isoflavone catabolism genes via mutagenesis study

To validate the involvement of the candidates in daidzein catabolism, we generated gene-disruption mutants of genes in the *ifc* cluster listed in Dataset S2. We found that, unlike the wild-type, V35 Δ*59390* was not able to degrade daidzein, indicating that locus 59390 encodes the initiating enzyme of the isoflavone catabolism pathway (Fig. 3a). In contrast, V35 Δ*59430* and the double mutant V35 Δ*59290* Δ*59540* exhibited distinctive peaks at Rt 1.7 and 1.5 min, respectively, compared to the wild-type (Fig. 3a), indicating that these genes are also involved in daidzein catabolism. Notably, V35 Δ*59430* accumulated a compound which was consistent with the compound X_1_ (Fig. S5). The Rt and mass spectrum of the compound X_1_ was identical to the authentic standard of 8-hydroxydaidzein (Fig. S6). We also found that the double mutant of locus 59540 and locus 59290, but not the single mutants of either gene, accumulated a distinctive peak at Rt 13.35 min, with a mass fragment ion at *m/z* 203.0353 (Fig. S7), suggesting that locus 59540 and locus 59290 are functionally redundant. The retention time and mass spectrum of the peak accumulated in V35 Δ*59290* Δ*59540* corresponded to the compound X_3_. Among the *ifc* catabolism intermediates, the compound X_3_ stably accumulated in the supernatant and remained after 48 h of cultivation. Using NMR analysis, we determined the chemical structure of compound X_3_ as 2-hydroxy-5-(4-hydroxyphenyl)-4*H*-pyran-4-one, a novel isoflavone catabolite (Fig. S8a). Gene disruption of locus 59470 did not affect the daidzein degradation ability. Collectively, even though the gene associated with the conversion of compound X_2_ to X_3_ were remained to be identified, we demonstrated that the genes located on locus 59390, 59430, 59540, and 59290 are isoflavone catabolism genes, thus named them *ifcA, B, D1,* and *D2*, respectively, and proposed an isoflavone catabolism pathway in *Variovorax* sp. (Fig. 2b).

**Fig. 3.**
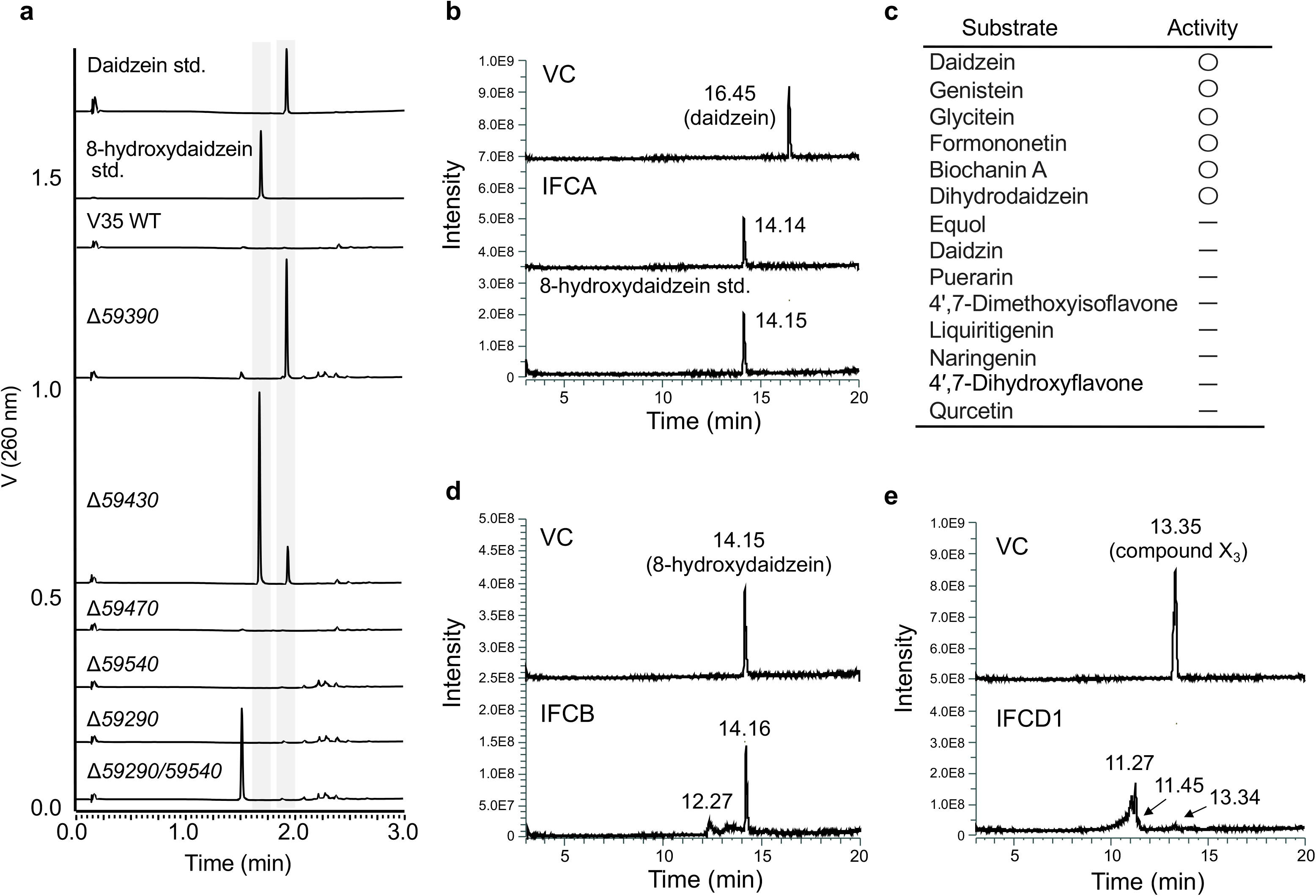
Daidzein degradation assay of *ifc* mutants and enzymatic activity of IFC enzymes. **(a)** UPLC analysis of culture supernatant of each *ifc* mutant grown in daidzein-supplemented MS medium. Essentially identical results were obtained in two independent experiments. **(b, d, and e)** LC-MS analysis of the reaction products from the recombinant proteins of IFCA, IFCB, and IFCD. A crude protein from *E. coli* transformed with an empty pET22b or pCold ProS2 vector was used as the negative control (VC). Total ion current chromatogram obtained in positive ionization mode with a full-scan range of *m/z* 50–350 is shown. Essentially identical results were obtained in two independent experiments. **(c)** Substrate specificity of IFCA. The catalytic activities of IFCA for each substrate is shown. Data were obtained from three technical replicates.

### Heterologous expression and functional characterization of IFC enzymes

We further characterized the catalytic activities of the recombinant IFC enzymes expressed in *E. coli*. When purified IFCA was incubated with daidzein in the presence of NADPH, a reaction product was detected. This product was identified as 8-hydroxydaidzein by direct comparison with the standard specimen (Fig. 3b and S9), indicating that IFCA catalyzes hydroxylation of the 8-position of daidzein. This reaction was dependent on the presence of NADPH (Fig. S10). We also examined the substrate specificity of IFCA using representative isoflavone aglycons, isoflavone glycosides, and other flavonoid families. IFCA accepted a series of isoflavones aglycons, including genistein, glycitein, formononetin, biochanin A, and dihydrodaidzein, with comparable catalytic efficiencies but did not give any products for 4′,7-dimethoxyisoflavone, equol, isoflavone glycoside, or any other flavones, flavanones, or flavonols (Fig. 3c and Fig. S11). This result suggests that IFCA is specific to isoflavone aglycons and that the methoxy moiety at the 7-position may inhibit substrate recognition and/or the catalytic activity of this enzyme. Given that IFCA formed a distinctive clade from other group A flavin-containing monooxygenases (FMOs) (supported by an 84.5% bootstrap portion: Fig. S12), these results indicated that IFCA represents a distinctive class of the group A FMOs that specifically catalyzes isoflavones.

Although a peak with a mass spectrum identical to the compound X_2_ was observed following the enzymatic reaction of recombinant IFCB protein with 8-hydroxydaidzein as a putative substrate (Fig. S13), 8-hydroxydaidzein remained after the reaction (Fig. 3d).

When IFCD1 was incubated with the compound X_3_ as a substrate, two peaks were detected with Rts of 11.27 and 11.45 min, respectively, and mass fragment ions were given at *m/z* 177.0555 and 193.0505, respectively (Fig. 3e and S14). Similarly, the reaction of IFCD2 with the compound X_3_ also gave the same two products (Fig. S15), further confirming the functional redundancy between IFCD1 and IFCD2. To further characterize the enzymatic reactions of IFCD1, as well as to estimate the chemical structure of the reaction products, we synthesized two stable isotopes of the compound X_3_, both of which are isotopically labeled at 3′ and 5′ or 2′, 3′, 5′, and 6′ positions, by biotransformation using the V35 Δ*ifcD1D2* mutant. These compounds were named X_3_-d_2_ and X_3_-d_4_, respectively (Fig. S16a and 17a). When IFCD1 was incubated with compound X_3_-d_2_ and X_3_-d_4_, the products had *m/z* values that were two and four mass units higher than those reacted with non-labeled compound X_3_, suggesting that compound X_3_ retained the B–ring of daidzein after reaction with IFCD1 (Fig. S16a and b, and S17a and b). The exact mass of the resulting compounds correlated with the calculated mass of C_10_H_10_O_3_ (177.0552) and C_10_H_10_O_4_ (193.0501), respectively. Thus, we predicted the chemical structures of these compounds (Fig. S18). In addition, functional annotation using KofamKOALA demonstrated that *Variovorax* isolates possessed gene sets involved in a meta-cleavage of catechol pathway, in which the aromatic ring of catechol is cleaved and eventually enter the tricarboxylic acid cycle, suggesting that daidzein is utilized for energy production.

### Distribution of *ifc* genes in bacterial kingdom

To assess the function of *ifc* genes in interactions with host legume plants, we established a *Variovorax* pan-genome by integrating 11 of our isolates and 83 publicly available genome sequences of *Variovorax* (Dataset S1b). Most of the *Variovorax* strains were isolated from soil or plant environments. *Ifc* genes were uniquely found in the strains isolated from plant-associated environments, especially from soybean and *Lotus japonicus* roots rich in isoflavones (Fig. 4). *Ifc* genes were also identified from strains isolated from plant species that do not synthesize isoflavones, such as maize, poplar, and *Avena barbata*. We further investigated the presence of *ifcA* genes in the bacterial kingdom by BLASTP search using IFCA as a query against the protein database at the National Center for Biotechnology Information (NCBI). IFCA homologs were uniquely identified from soil bacteria belonging to phylum Proteobacteria, including *Variovorax*, *Burkholderia*, *Paraburkholderia*, *Bradyrhizobium*, *Pseudomonas*, and *Sphingomonas*, but not found in intestinal bacteria species, and the *ifc* cluster including *ifcA/B/D1/D2* was conserved in those strains (Fig. S20). Phylogenic analysis of IFCA further demonstrated that IFCA was divided into two distinctive groups. Even though further functional characterization is needed, the two IFCA groups may have functional differences, such as substrate specificity.

**Fig. 4.**
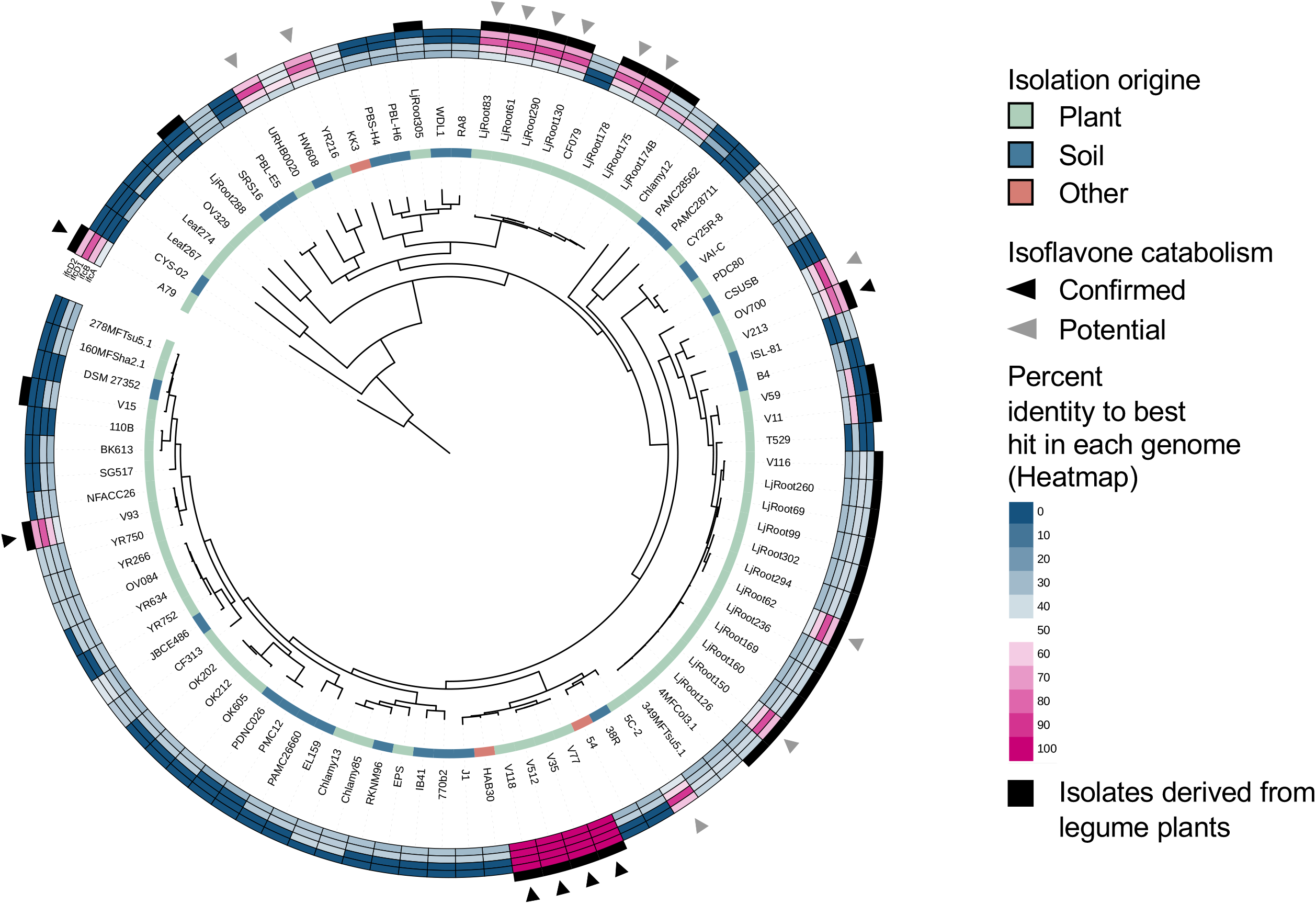
Phylogenetic distribution of *ifc* genes in genus *Variovorax*. The phylogenetic tree of 92*Variovorax* genomes was inferred from aligned single-copy genes using the MLE method. The inner ring depicts the isolation source of each genome. Heatmap representing per cent identity of BLASTP hits in each *Variovorax* genome to *ifcA/B/D1/D2* from strain V35. The black outer rings represent the strains isolated from legume plants. Arrowheads indicate the isoflavone-degrading strains. Black color indicates the isoflavone degrading strains whose degradation abilities were experimentally confirmed. Gray color indicates the potential isoflavone-degrading strains predicted by *in silico* analysis.

To test whether *ifcA* gene is involved in the bacterial adaptation to the isoflavone containing environment, we measured the *in vitro* bacterial growth in daidzein-containing nutrient medium. We found that the growth of the V35 Δ*ifcA* mutant was significantly inhibited in the presence of daidzein, whereas no clear growth inhibition was observed by daidzein supplementation in the wild-type strain (Fig. 5). Collectively, our findings imply that *ifc* contributes to the bacterial adaptation to isoflavone-rich environments such as legume roots.

**Fig. 5.**
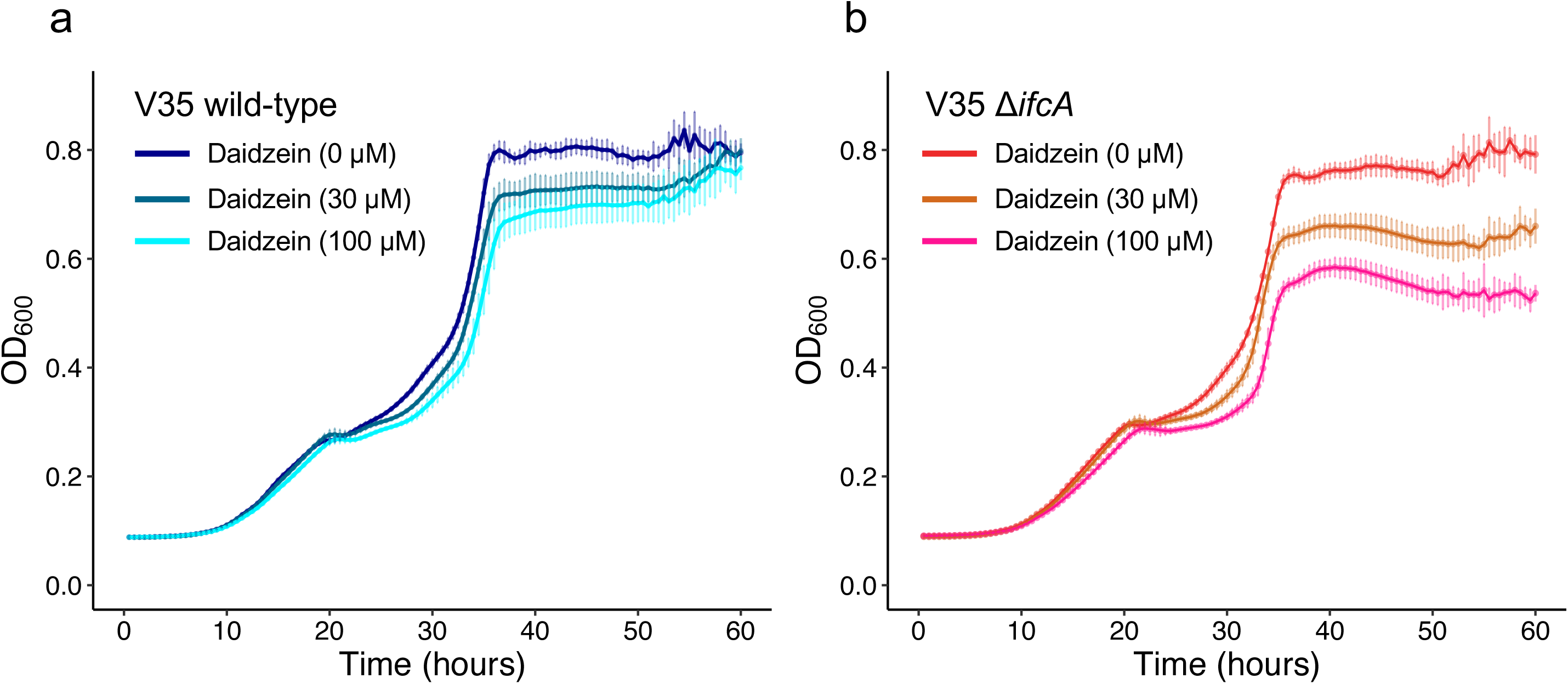
*In vitro* growth assay of V35 and *ifcA*-deficient mutant. Growth of V35 wild-type **(a)** and V35 Δ*ifcA* mutant **(b)** in peptone-beef extract medium and peptone-beef extract medium supplemented with 30 or 100 μM daidzein. Essentially identical results were obtained from three independent experiments. Data are expressed as the mean ±SD (n = 6).

## Discussion

The root microbiota is critical for sustainable agricultural production due to its fundamental role in plant growth and health (37–39). Unveiling the metabolic pathway of PSMs in the microbiota is a prerequisite for rational design to optimize the function of root microbiota for agricultural purposes. Here, we isolated isoflavone-degrading soil bacteria from soybean roots and discovered an oxidative catabolism pathway of isoflavone, which is distinct from the reductive isoflavone catabolism found in human intestinal bacteria optimized for anaerobic gut environments (40).

Several flavonoid catabolism pathways have been identified from soil bacterial species. Quercetinases identified from *Streptomyces* sp. FLA, and *Bacillus subtilis* YxaG, for example, catalyze the 2,4-dioxygenolytic cleavage of quercetin (41, 42). *Pseudomonas putida* PML2 catabolizes quercetin by C-ring cleavage starting from 3,3′-didehydroxylation (43). While flavonoid catabolism in these bacteria species is initiated by C-ring cleavage, *fde* genes in *Herbaspirillum seropedicae* SmR1 encode enzymes that catabolize naringenin by 8-hydroxylation and subsequent dioxygenation of the A-ring (34). These findings illustrate that bacterial oxidative flavonoid catabolism is distinctively divided into either A- or C-ring cleaving pathways.

The Isoflavone catabolism pathway of *Variovorax* revealed in this study resembles the A-ring cleaving pathway defined by the *fde* genes. In contrast, it has been reported that *Rhizobium* sp. strain NGR234 degraded daidzein via C-ring cleavage fission (29), implying another oxidative isoflavone catabolism pathway initiated from C-ring cleavage in soil bacteria.

We also identified several catabolites in the isoflavone catabolism of a *Variovorax* species, including a previously unidentified compound, 2-hydroxy-5-(4-hydroxyphenyl)-4*H*-pyran-4-one (Fig. 2b). Among the identified catabolites, 8-hydroxydaidzein, an enzymatic reaction product of IFCA, has been found only in fermented soy food and exhibits a series of beneficial activities for human health, such as antioxidant, antitumor, antimelanogenesis, aldose reductase inhibitory, and hepatoprotective activities (44). Therefore, there have been many attempts to produce 8-hydroxydaidzein by biotransformation and to discover the novel enzymes that efficiently catalyze the hydroxylation of the A-ring of isoflavone (45). Given the impact of isoflavone hydroxylation in the food industry and therapeutics, our findings provide valuable insights into isoflavone biotransformation and contribute to the exploration of novel bioactive isoflavone derivatives.

The PSM catabolic abelites of plant-associated microorganisms play a crucial role in their interaction with host plants and in the formation of plant species-specific root microbiota. Comparative genome analysis revealed that *Variovorax* strains isolated from the roots of legume plants such as soybean and *L. japonicus* often possess *ifc* genes (Fig. 4). This observation is in line with our previous report that *Arthrobacter* isolates-derived from tobacco roots harbored a characteristic genomic feature, enabling them to utilize tobacco-specific PSMs such as nicotine and santhopine (11). PSM catabolic abilities may contribute to predominant colonization in the roots and rhizosphere, where PSMs in general inhibit the growth of microorganisms (46–48). In fact, disruption of the *ifcA* gene reduced the growth of V35 strains in media containing daidzein (Fig. 5). Combined with the fact that isoflavone modulates the taxonomic structures of soybean root microbiota (19), *ifc* genes might underly the isoflavone-mediated assemblage of legume root microbiota.

Our comparative genome analysis revealed that *ifc* genes are distributed in phylum Proteobacteria, including *Bradyrhizobium* and *Burkholderia* species, which are known to establish a symbiotic relationship with legume plants (49, 50). Interestingly, *Burkholderia* strains possessing *ifc* genes, such as *Burkholderia* sp. USMB20 and *Paraburkholderia steynii*, have been isolated from legume nodules (51, 52), suggesting the association between *ifc* genes and symbiotic interactions with legume plants. In fact, it has been reported that flavonoid metabolites produced by symbiotic rhizobia affect *nod* gene expression and bacterial growth. Chalcones, a luteolin degradation intermediate of *Rhizobium meliloti* (29), exhibits stronger induction activity of *nod* genes compared to luteolin (53). *Bradyrhizobium* sp. strain ORS285 converted naringenin into its *O*-methylated form, which stimulates the growth of strain ORS285 (54). Collectively, our findings set a new trajectory toward a molecular understanding of plant-microbiota interactions and their potential application in sustainable agriculture.

## Materials and methods

### Bacterial strains and chemicals

The bacterial strains and plasmids used in this study are listed in Dataset S2. The bacterial strains belonging to Comamonadaceae were isolated from soybean roots grown in nitrogen-deficient conditions. A detailed description and isolation procedures can be found in Yazaki *et al.* (32). All *Variovorax* isolates were grown at 28°C in peptone-beef extract medium (10 g/L peptone, 10 g/L beef extract, and 5 g/L NaCl) supplemented with appropriate antibiotics (5 μg/mL tetracycline). *Escherichia coli* strains were grown at 37°C in Luria–Bertani (LB) medium supplemented with appropriate antibiotics (50 μg/mL kanamycin and 5 μg/mL tetracycline). Chemicals were obtained from Wako Pure Chemical Industries (Japan) or Nacalai Tesque (Japan), otherwise listed in Dataset S2.

### Plasmid construction and generation of gene knockout mutants

All constructions and primers used in this study are listed in Dataset S2. Since strain V35 exhibits kanamycin resistance, pT18*mobsacB* was generated from pK18*mobsacB* by replacing the *npt II* gene with the tetracycline resistance gene. The tetracycline resistance gene was artificially synthesized with *Bgl2* and *Nco1* recognition sequences (Genewiz, United States) and were cloned into *Bgl2* and *Nco1* sites of pK18*mobsacB* to form pT18*mobsacB*.

The internal fragments of the target genes were amplified by KOD FX Neo (Toyobo, Japan) and cloned into pT18*mobsacB* by in-fusion reaction (Clontech, United States) for the construction of insertion mutants. For the construction of deletion mutants, the upstream and downstream flanking regions of the target genes were amplified and cloned into pT18*mobsacB* by in-fusion reaction. The constructed vectors were introduced into V35 by triparental mating*. E. coli* DH5α containing the constructed plasmid was used as a donor strain and *E. coli* HB101 harboring pRK2013 was used as a helper strain (Figurski and Helinski 1979). *E. coli* strains were grown in LB medium supplemented appropriate antibiotics, and V35 was grown in peptone-beef extract medium. Each bacterial strain was mixed and washed three times with peptone-beef extract medium, then spotted onto 0.45-μm MCE membranes (Merck Millipore, United States) placed on peptone-beef extract medium agar plates. After drying, plates were incubated at 28°C overnight. An initial recombinant was selected on peptone-beef extract medium supplemented with 10 μg/mL of tetracycline and 100 μg/mL of ampicillin. The obtained colonies were further cultured overnight at 28°C in 2 mL of peptone-beef extract medium to induce a second recombination event, and the deletion mutant was selected on peptone-beef extract medium supplemented with 10% sucrose. The constructed mutants were screened for antibiotic resistance and confirmed by PCR.

### Measurements of isoflavone-degrading activities using resting cells

Bacterial strains were cultured overnight at 28°C in 2 mL of peptone-beef extract medium. Bacterial cells were collected by centrifugation at 1,500 x *g* for 10 min and washed twice with mineral salt (MS) medium (55). Bacterial pellets were resuspended in 2 mL of MS medium supplemented with daidzein at a concentration of 4 μM and incubated for 3 days. After centrifugation at 1,500 x *g* for 10 min, supernatants were collected and stored at –80°C until use. To extract isoflavones from the supernatants, 500 μL of supernatant was acidified with 5 μL of 1 M HCl and mixed with 500 μL of ethyl acetate. After centrifugation at 9,100 x *g* for 10 min, 400 μL of organic layer was collected and evaporated. The extracts were resuspended in 95% MeOH containing 0.1% (v/v) formic acid. After filtering through a 0.45-μm Minisart RC4 filter (Sartorius, Germany), collected samples were stored at –30°C until ultra-high-performance liquid chromatography (UPLC) analysis.

### Time course daidzein degradation activity of strain V35

Bacterial strains were cultured overnight at 28°C in 2 mL of peptone-beef extract medium. Bacterial cells adjusted to OD_600_ = 1.0 were collected by centrifugation at 13,000 x *g* for 1 min and washed twice with MS medium. The resulting bacteria pellet was resuspended in 2 mL of MS medium supplemented with daidzein a concentration of 20 μM and cultured at 28°C. For the time course daidzein degradation assay of the V35 wild-type strain, 100 μL of culture supernatant was collected at 0, 1, 2, 4, and 8 h, immediately mixed with equivalent volume of methanol, and centrifuged at 13,000 x *g* for 1 min. Daidzein degradation of *ifc* mutants was conducted for 48 h of cultivation. After filtration through a 0.45-μm Minisart RC4 filter (Sartorius, Germany), collected samples were stored at –30°C until UPLC and LC-MS analysis.

### Heterologous expression of recombinant proteins in *E. coli*

Target genes were amplified by KOD FX Neo (Toyobo, Japan). The amplified *ifcD1* sequence was cloned into the pET-22b(+) (Merck KGaA, Germany) and the amplified *ifcA*, *ifcB*, and *ifcD2* sequences were cloned into the pCold ProS2 by in-fusion reaction. The used primer sets, vector and, *E. coli* host are summarized in Dataset S2. *E. coli* strain BL21 (DE3) transformed with the constructed vector was grown at 37°C in LB medium supplemented with 100 μg/mL of ampicillin until its OD_600_ reached 0.5. Recombinant protein expression was induced by adding isopropyl β-D-1-thiogalactopyranoside a final concentration of 0.5 mM. *E. coli* strains harboring pET-22b(+) were cultured for 22 h at 18°C, and strains harboring pCold ProS2 were cultured 24 h at 15°C. The cell pellets were washed two times with cold Tris-HCl buffer (150 mM NaCl, 10 mM Tris-HCl, pH8.0), divided into four equal parts, and stored at −30°C until *in vitro* assay.

Each cell pellet was resuspended in 1 mL of cold lysis buffer (50 mM sodium phosphate (pH 8.0), 300 mM NaCl, and 10 mM imidazole), and sonicated four times for 30 sec, with 30-sec intervals, using an ultrasonic homogenizer Sonifier Model 250A (Branson, United States) with a duty cycle of 40% and an output control of 20%. The homogenate was centrifuged at 13,000 x *g* for 5 min at 4°C, and the supernatant was applied to Ni-NTA chromatography. One hundred fifty microliters of Ni-NTA agarose (QIAGEN, Germany) was loaded into Micro Bio-Spin Chromatography Column (0.8 mL bed volume) (Bio-Rad, United States). The resin was equilibrated with 600 μL of the lysis buffer and centrifuged at 1,000 x *g* for 1 min at 4°C to remove buffer. His-tagged proteins in 600 μL of the supernatant were loaded and passed through the column by centrifugation at 1,000 x *g* for 1 min at 4°C. They were then washed twice with 600 μL of a wash buffer (50 mM sodium phosphate (pH 8.0), 300 mM NaCl, and 20 mM imidazole). The adsorbed proteins were eluted twice with 100 μL of an elution buffer (50 mM sodium phosphate (pH 8.0), 300 mM NaCl, and 250 mM imidazole). The expression level of soluble protein was confirmed by SDS-PAGE, and protein concentration was measured using a Qubit Protein Assay Kit (Thermo Fisher Scientific, United States).

### Substrate preparation for IFCD

The V35 Δ*ifcD1D2* was cultured at 28°C in 2 mL of peptone-beef extract medium supplemented with 5 μg/mL tetracycline as pre-culture, then scaled up to 100 mL of peptone-beef extract medium and cultured for 24 h at 28°C. Bacterial cells adjusted to OD_600_ = 1.0 were collected by centrifugation at 13,000 x *g* for 1 min and washed twice with MS medium. The resulting bacteria pellet was resuspended in 100 mL of MS medium supplemented with daidzein at a final concentration of 33 µg/mL. After 3 days of cultivation at 28°C, bacterial cells were collected by centrifugation at 13,000 x *g* for 5 min. Culture supernatant was adjusted to pH 2.0 with hydrochloric acid. After an equivalent volume of ethyl acetate was added, the organic layer was collected and evaporated. The precipitate was resuspended in 1 mL of MeOH using sonication. After filtration with a 0.45-μm Minisart RC4 filter (Sartorius, Germany), MeOH was dried under nitrogen gas. The resulting precipitate was adjusted to 1 μM with DMSO.

For NMR analysis, compound X_3_ was purified using an HPLC system (Shimadzu, Japan) equipped with a Develosil C30-UG-3 column (20 × 150 mm, NOMURA CHEMICAL). The mobile phase was 30% acetonitrile with 0.3% acetic acid. The effluent was monitored at 260 nm using photo diode array (SPD-M10A, Shimadzu, Japan).

For isotopically labeled daidzein, V35 Δ*ifcD1D2* was cultured for 3 days at 28°C in 2 mL of MS medium supplemented with daidzein-3’,5’,8-d3 or daidzein-2’,3’,5’,6,6’,8-d6 at final concentrations of 5 µg/mL. Bacterial cells were collected by centrifugation at 4,000 rpm for 10 min, then mixed with 40 μL of 1M hydrochloric acid and equivalent volume of ethyl acetate. After centrifugation at 13,000 x *g* for 1 min, the organic layer was collected and dried under nitrogen gas. The resulting precipitate was adjusted to 10 mM with DMSO.

### NMR analysis of compound X_3_

The purified compound X_3_ was analyzed by NMR (500 MHz, Avance 500, Bruker, Germany) and confirmed by proton nuclear magnetic resonance (1H-NMR) and 1H–1H double quantum filtered chemical shift correlation spectroscopy (DQF-COSY). The chemical shifts were assigned relative to the solvent signal. DMSO-d6 was used as the solvent. [^1^H NMR (DMSO-d_6_): 7.66 (1H, *s*, H-6), 7.19 (2H, *d*, *J* = 8.8 Hz, H-2’, H-6’), 6.76 (2H, *d*, *J* = 8.6 Hz, H-3’, H-5’), 5.47 (1H, *s*, H-3)] (Fig. S8 b).

### Measurement of enzymatic activity of recombinant proteins

An *in vitro* assay of recombinant IFCA was performed using 100 μL of a reaction mixture consisting of 50 mM sodium phosphate (pH 7.5), 1 mM NADPH used as a coenzyme, 100 μM substrates, and 5 μg of each recombinant protein. The enzymatic activities of IFCB were performed using 100 μL of a reaction mixture consisting of 50 mM sodium phosphate (pH 7.5), 100 μM substrate, and 5 μg of each recombinant protein. The enzymatic activities of IFCD were measured with 50 μM of each of the prepared substrate in the reaction mixtures used for IFCB. All the reactions were conducted at 28°C for 30 min and terminated by the addition of 100 μL of MeOH. After filtration through a 0.45-μm Minisart RC4 filter (Sartorius, Germany), the resulting reaction mixtures were subjected to UPLC and LC-MS analysis.

### UPLC and LCMS analysis of reaction products

The Microbial and enzymatic reaction products were analyzed by UPLC (Shimadzu, Japan). For each sample, 5 μL was injected onto an Acquity UPLC BEH C18 column (1.7 μm, 2.1 × 50 mm^2^; Waters, United States) with a UPLC BEH C18 VanGuard Precolumn (1.7 μm, 2.1 × 5 mm^2^, Waters, United States). The column oven temperature was set at 40°C. The LC mobile phases consisted of water (mobile phase A) and acetonitrile (mobile phase B), each containing 0.3% (v/v) formic acid. Elution was set at a rate of 1.0 mL/min with an isocratic elution of 5% B over 0.5 min, followed by a linear gradient from 5% to 80% B over 2.5 min.

LC-MS analysis was conducted with Orbitrap LC-MS (Thermo Fisher Scientific, United States). For each sample, 5 μL was injected onto a Shim-pack Velox Biphenyl column (1.8 μm, 2.1 × 100 mm^2^; Shimadzu, Japan) with a Shim-pack Velox EXP Guard Column Cartridge (Biphenyl UHPLC, 2.1 × 5 mm^2^; Shimadzu, Japan). The column oven temperature was set at 40°C. The mobile phases were water containing 0.1% (v/v) formic acid (solvent A) and methanol containing 0.1% (v/v) formic acid (solvent B). Elution was set at 0.3 mL/min with an isocratic elution of 0% B over 5 min, followed by a linear gradient from 0% to 90% B over 15 min. The mass spectra were obtained in the negative electrospray ionization mode with following settings: ionization, ESI; spray voltage, + 2.0 kV; capillary temperature, 300°C; sheath gas flow rate, 55 L/min; aux gas flow rate, 15 L/min; sweep gas flow rate, 3 L/min; aux gas heater temperature, 450°C; S-lens RF level 50.0; collision energy, 15 eV. The MS scan was in the range of *m/z* 50–350. Data acquisition and analysis were performed using FreeStyle 1.1 SP1 (Thermo Fisher Scientific, United States).

### RNA extraction and RNA-seq analysis

Strain V35 was precultured in 3 mL of peptone-beef extract medium at 28°C overnight and scaled up to 20 mL of peptone-beef extract medium. After overnight cultivation at 28°C, bacterial cells were collected by centrifugation at 5,800 x *g* rpm for 10 min and washed twice with MS medium. The resulting bacterial pellet was resuspended in MS medium or MS medium supplemented with daidzein at final concentration of 30 μg/mL and cultured for 16 h at 28°C. One hundred microliters of bacterial culture was harvested at 15,300 x *g* rpm for 2 min, immediately frozen in liquid nitrogen, and stored at –80°C until RNA extraction.

Total RNA was extracted using TRI Reagent (Cosmo Bio Co., Ltd., Japan) according to the manufacturer’s instructions and purified by phenol–chloroform extraction. The quality and concentration of the extracted RNA were evaluated using a BioSpec-nano (Shimadzu, Japan) and Qubit Quantification Platform RNA BR Assay Kit (Invitrogen, United States). The extracted RNA was sent to Macrogen (Korea) for library preparation using the TruSeq stranded mRNA Library (Illumina, United States) and sequencing of 2 × 100 bp paired-end reads using the NovaSeq 6000 (Illumina, United States). Obtained raw reads were mapped to strain the V35 genome using Kallist 0.43.1 with the default setting (Bray et al. 2016). The raw sequence reads were submitted to the DNA Data Bank of Japan (DDBJ; PRJDB16019) to be publicly available.

### Whole genome sequencing and comparative genome analysis

The genome DNA extraction, whole-genome sequencing, and subsequent assembly were performed as described previously (11). In brief, the cells were lysed by lysozyme and proteinase K, and the resulting total DNA was extracted with phenol and chloroform. Each genomic DNA was subjected to 20-kbp fragmentation using the Megaruptor2 (Diagenode, Belgium). Subsequently, SMRTbell Express Template Prep Kit 2.0 (Pacific Biosciences, United States) was employed to construct the library. To distinguish between different fragments, barcodes were attached to each fragmented genome. The resulting samples were then pooled and size-selected using the BluePippin system (Sage Science, United States) with a cutoff at 15-kbp. Sequencing of the genomic library was conducted on a single PacBio Sequel II system 2.0 cell. The obtained reads were assembled using HGAP4 via SMRT link (v 8.0.0). The sequence dataset was submitted to the DDBJ (PRJDB16018) to be publicly available.

Public genome sequences of *Variovorax* were retrieved from the Integrated Microbial Genomes & Microbiomes system (https://img.jgi.doe.gov) and the NCBI (https://www.ncbi.nlm.nih.gov). Using a total of 92 *Variovorax* genomes, putative protein-coding sequences were predicted using Prokka (56). The annotation of candidate ORFs was then conducted using DFAST (57) . High-resolution phylogenetic inference was conducted using Orthofinder2 (58). The identification of homologous *ifc* genes was performed by BLASTP using amino acid sequences of IFCA/B/D1/D2 against the proteome predicted by the *Variovorax* genome sequences (E-value thread > 1 × 10^−15^). Phylogenetic trees and associated data were visualized using iTOL version 5 (59). KEGG pathway analysis of V35 was performed using KofamKOALA (60)

### *In vitro* bacterial growth assay

V35 and V35 Δ*ifcA* was cultured in 3 mL of peptone-beef extract medium for 16 h at 28°C with shaking at 180 rpm. The overnight cultures were diluted in peptone-beef extract medium and adjusted to OD_600_ = 0.001. Different concentrations of daidzein were added to the bacterial cultures, and applied into a 96-well microplate. Growth curves were monitored under the wavelength of OD_600_ nm with an interval of 30 min at 28°C, by Synergy HTX (BioTek Instruments, United States).

## Acknowledgments

We thank Ms. Keiko Kanai and Ms. Rie Mizuno for technical assistance; We would like to thank Dr. Shin Okazaki, Dr. Safirah Tasa Nerves Ratu, Dr. Ryohei Thomas Nakano, Dr. Hiroaki Kusano, and Dr. Ryosuke Munakata for helpful discussion. We would like to thank DASH/FBAS, Research Institute for Sustainable Humanosphere, Kyoto University, for supporting institutional setting. This study was supported in part by grants from the Core Research for Evolutional Science and Technology, Japan Science and Technology Agency (CREST, JST; JPMJCR17O2 to Y.A. and A.S.); Japan Society for the Promotion of Science KAKENHI (21H02329 to A.S., 20H05592 to S.M., and 22H00364 to K.S.); and the Research Institute for Sustainable Humanosphere (Mission 1) to A.S.

T.S. and A.S. conceived and designed the research. A.A., S.K., O.J., K.Y., and A.S. supervised the experiments. W.Y. and T.S isolated bacteria strains and performed RNA-seq experiment. T.S., W.Y., S.M., A.S., and K.S. conducted whole-genome sequencing. W.Y., T.S., and N.A. analyzed transcriptome and bacterial genome data. N.A. and T.S. generated *ifc* gene disrupted mutants and measured isoflavone degradation ability. N.A., T.S., M.N. and A.S. measured of the enzymatic activities using resting cells and recombinant proteins. S.K. conducted NMR analysis. T.S., N.A., K.S., and A.S. wrote the article with contributions of all the authors. A.S. agrees to serve as the author responsible for contact and ensures communication.

## Data availability

The data set supporting the results of this study is publicly available at the DDBJ (https://www.ddbj.nig.ac.jp) (PRJDB16018 and PRJDB16019). Additional data related to this paper will be made available from the corresponding author upon reasonable request.

## Supplementary materials

**Fig. S1. Daidzein degradation assay of soybean-root associated bacteria isolates.** UPLC analysis of culture supernatant of each bacterial strain belonging to Comamonadaceae grown in MS medium supplemented with daidzein. Essentially identical results were obtained in two independent experiments.

**Fig. S2. LC-MS/MS analysis of daidzein catabolism intermediates in *Variovorax* sp. strain V35. (a–c)** Product ion spectra of compound X_1_-X_3_. Essentially identical results were obtained from two independent experiments.

**Fig. S3. Time-course daidzein degradation activities of daidzein- and mock-treated strain V35.** Residual substrate contents in reaction mixture are shown relative to that at the start of the reaction. Essentially identical results were obtained from two independent experiments. Data are expressed as the mean ±SD (n = 3).

**Fig. S4. Comparative analysis of *ifc* genes with *fde* operon and proposed naringenin catabolism pathway by *fde* genes. (a)** Synteny analysis and visualization between *ifc* genes and *fde* operon were performed using clinker (61). Per cent identity of shared genes between *ifc* and *fde* gene clusters are represented in greyscale. The functional assignments and per cent identity of each *fde* gene to the *ifc* genes from strain V35 are shown in the table below the synteny map. **(b)** Proposed naringenin catabolism pathway by *fde* genes in *Herbaspirillum seropedicae* strain SmR1 (34).

**Fig. S5. LC-MS analysis of daidzein degradation products of** V35 Δ*59390*. Total ion chromatogram negative ionization mode with full-scan range of *m/z* 50– 350, mass and product ion spectra for culture supernatant from V35 Δ*59390*.

**Fig. S6. LC-MS analysis of authentic standard of 8-hydroxydaidzein.** Total ion chromatogram negative ionization mode with full-scan range of *m/z* 50–350, mass and product ion spectra for authentic standard of 8-hydroxydaidzein.

**Fig. S7. LC-MS analysis of daidzein degradation products of** V35 Δ*59290* Δ*59540*. Total ion chromatogram negative ionization mode with full-scan range of *m/z* 50-350, mass and product ion spectra for culture supernatant from V35 Δ*9290* Δ*59540*.

Fig. S8. NMR analysis of compound X_3_

(a) Chemical structure of compound X_3_ for NMR analysis and (b) 1H NMR spectrum.

**Fig. S9. NADPH requirement for catalytic activity of IFCA.** UPLC analysis of reaction product using the recombinant protein of IFCA without NADPH and with NADPH, and authentic standard of 8-hydroxydaidzein. Essentially identical results were obtained in two independent experiments with three technical replicates.

**Fig. S10. LC-MS analysis of the reaction product using the recombinant protein of IFCA and authentic standard of 8-hydroxydaidzein.** The mass and product ion spectra for **(a)** reaction product from the recombinant protein of IFCA and **(b)** authentic standard of 8-hydroxydaidzein. Essentially identical results were obtained in three independent experiments with three technical replicates.

Fig. S11. Substrates specificity of IFCA.

**Fig. S12. Phylogenetic analysis of group A flavin-containing monooxygenase.** The phylogenic tree of group A flavin-containing monooxygenase was constructed by MLE method referring to Westphal et al. Bootstrap values (1,000 replicates) above 90% are indicated by empty circles on nodes. Some branches forming big clades were collapsed, giving the representative enzyme designation. 2,6-dihydroxypyridine-3-monooxygenase is from *P. nicotinovorans* (DhpH) (UniProtKB: Q93NG3). FAD-dependent urate hydroxylase from *K. pneumoniae* (HpxO) (UniProtKB: B6D1N4). FAD-dependent monooxygenase from *P. aeruginosa* (PqsL) (UniProtKB: Q9HWJ1). Monooxygenase from *S. venezuelae* (JadH) (UniProtKB: Q5U913). Epoxidase from *S. lasalocidi* (LasC) (UniProtKB: B5M9L6). 3-(3-hydroxy-phenyl)propionate/3-hydroxycinnamic acid hydroxylase is from *C. testosteroni* (MhpA) (UniProtKB: Q9S158). FAD-dependent monooxygenase is from *E. nidulans* (AfoD) (UniProtKB: Q5BEJ7). *Maackiain detoxification* from *F. solani* (MAK1) (UniProtKB: Q01446).

**Fig. S13. LC-MS analysis of the reaction product from the recombinant protein of IFCB and compound X_2_.** The mass and product ion spectra for **(a)** reaction product from the recombinant proteins of IFCB and **(b)** compound X_2_. These data are representative of two independent experiments.

**Fig. S14. LC-MS analysis of the reaction products using the recombinant protein of IFCD1.** The mass and product ion spectra for reaction product **(a)** 1 and **(b)** 2 using the recombinant proteins of IFCD1. These data are representative of two independent experiments.

**Fig. S15. LC-MS analysis of the reaction products using the recombinant protein of IFCD2.** The mass and product ion spectra for reaction product **(a)** 1 and **(b)** 2 using the recombinant proteins of IFCD2. These data are representative of two independent experiments.

**Fig. S16. Chemical structure of Compound X_3_-d_2_ and LC-MS analysis of reaction products with the recombinant protein of IFCD1. (a)** Chemical structure of Compound X_3_-d_2_. The mass and product ion spectra for reaction product **(b)** 1 and **(c)** 2 using the recombinant protein of IFCD1. These data are representative of two independent experiments.

**Fig. S17. Chemical structure of Compound X_3_-d_4_ and LC-MS analysis of reaction products with the recombinant protein of IFCD1. (a)** Chemical structure of Compound X_3_-d_4_. The mass and product ion spectra for reaction product **(b)** 1 and **(c)** 2 using the recombinant protein of IFCD1. These data are representative of two independent experiments.

**Fig. S18.** Predicted the chemical structure of reaction product using the recombinant protein of IFCD. **(a)** C_10_H_10_O_3_ and **(b)** C_10_H_10_O_4_.

**Fig. S19. KEGG pathway map for meta-cleavage of catechol pathway (M00569) in *Variovorax* sp. V35**. Green arrows indicated conserved pathway in *Variovorax* sp. V35.

**Fig. S20. Phylogenetic analysis of IFCA homologues protein and comparative analysis of *ifc* gene cluster. (a)** The phylogenic tree of IFCA homologues protein was constructed by MLE method. The inner ring depicts the isolation source of each genome. The bar on the right denotes the family classification of each strain. Two distinctive IFCA groups are highlighted by blue and red ranges. **(b)** Synteny analysis of *ifc* cluster between taxonomically distinct bacterial species. Per cent identity of shared genes between *ifc* and *fde* gene cluster are represented in greyscale.

