## Supplemental Materials for "Discovery of an isoflavone oxidative catabolic pathway in legume root microbiota"

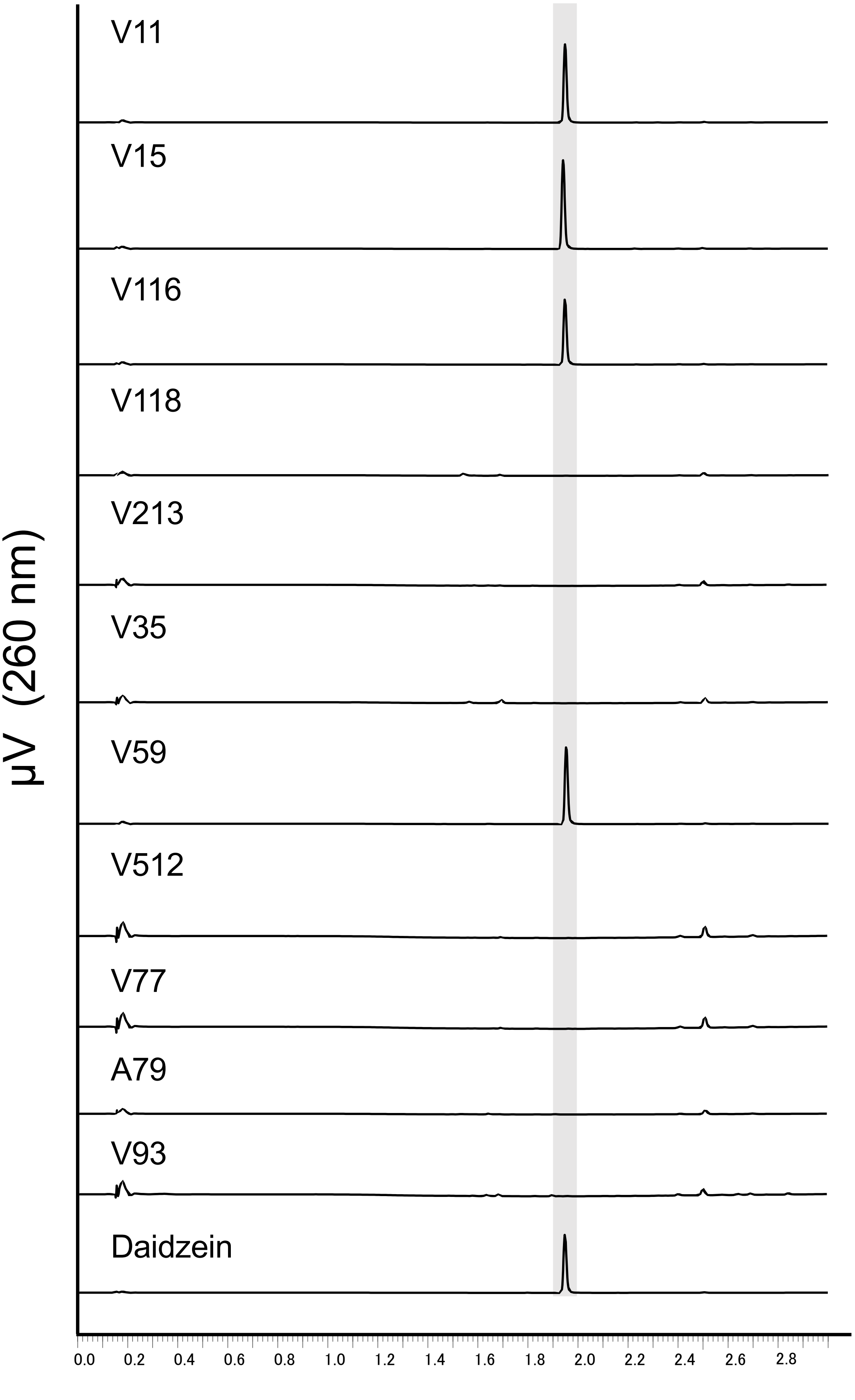


Fig. S1

Aoki et al.

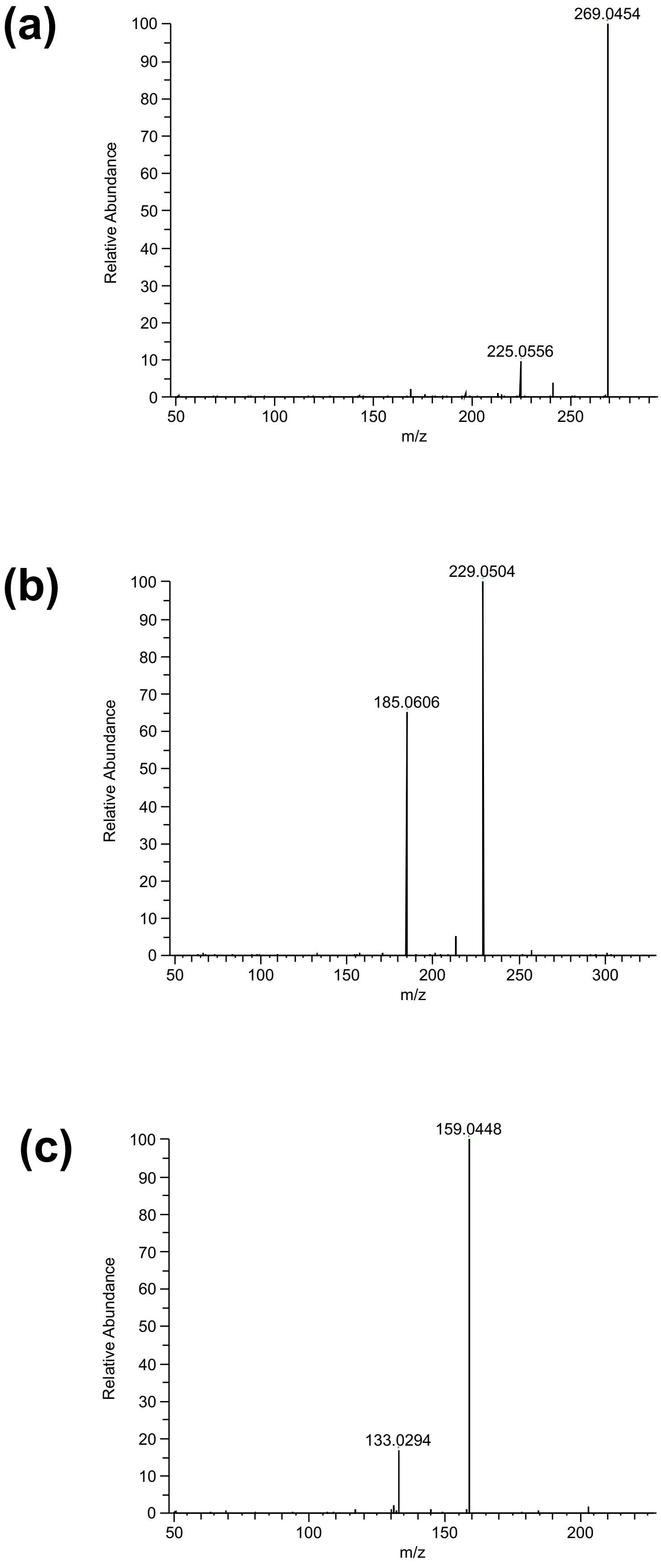


Fig. S2 Aoki et al.

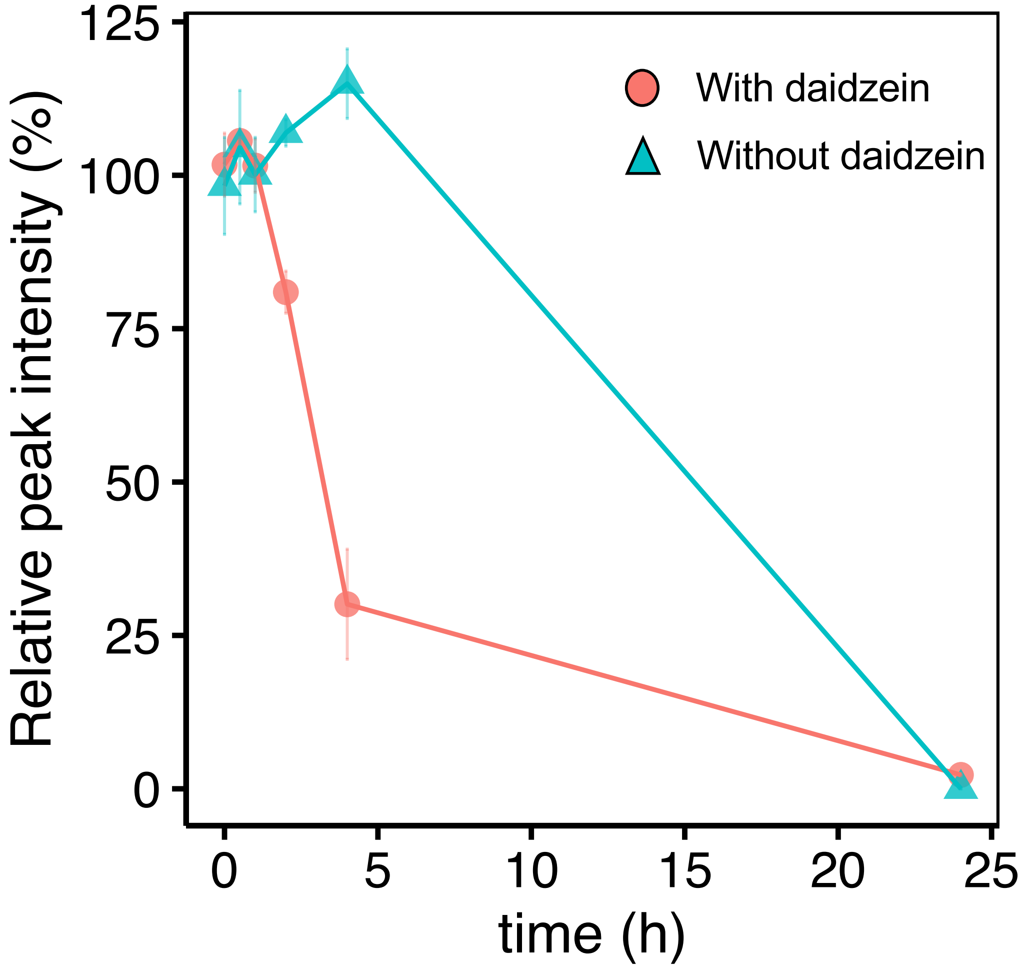


Fig. S3

Aoki et al.

**(a)**

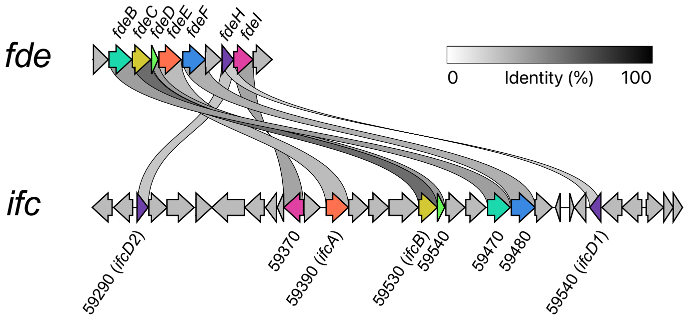


| Gene | Annotation | Homologus gene (% identity) |
|---|---|---|
| *fdeA* | Protein involved in meta-pathway of phenol degradation | not hit |
| *fdeB* | alpha/beta hydrolase | 59470 (48%) |
| *fdeC* | glyoxalase | 59430 (66%) |
| *fdeD* | (2Fe-2S)-binding protein | 59440 (55%) |
| *fdeE* | FAD-dependent 2-polyprenyl-6-methoxyphenol hydroxylase protein | 59390 (36%) |
| *fdeF* | MFS transporter | 59480 (45%) |
| *fdeG* | Cyclase | not hit |
| *fdeH* | cupin | 59290 (37%)/ 59540 (35%) |
| *fdeI* | 2-keto-4-pentenoate hydratase | 59370 (45%) |
| *fdeJ* | UDP-glucose 4-epimerase | not hit |

**(b)**

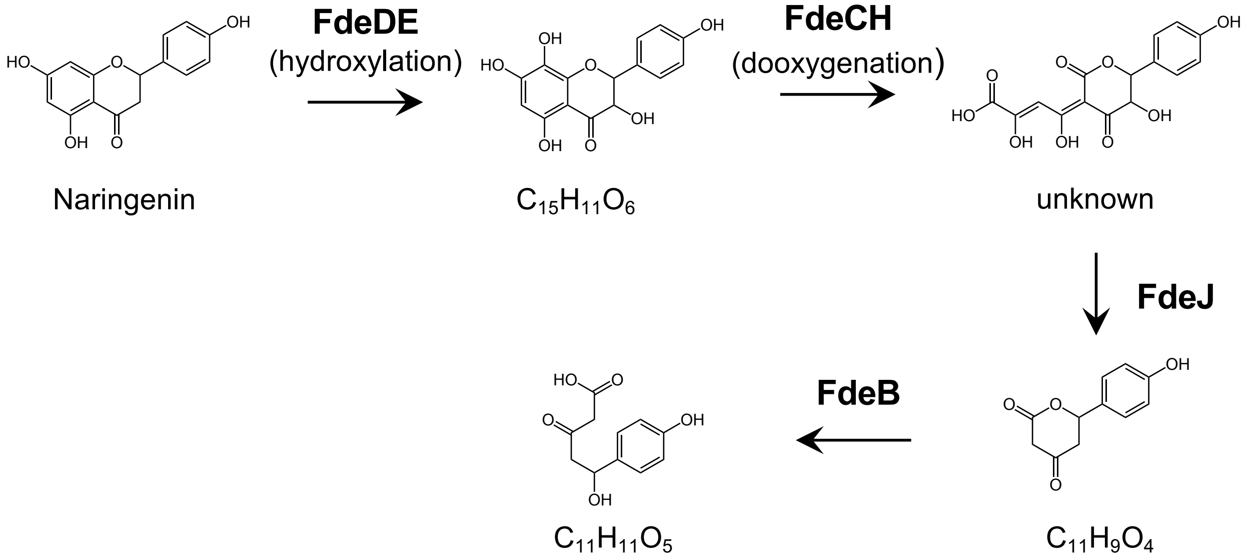


Fig. S4

Aoki et al.

TIC

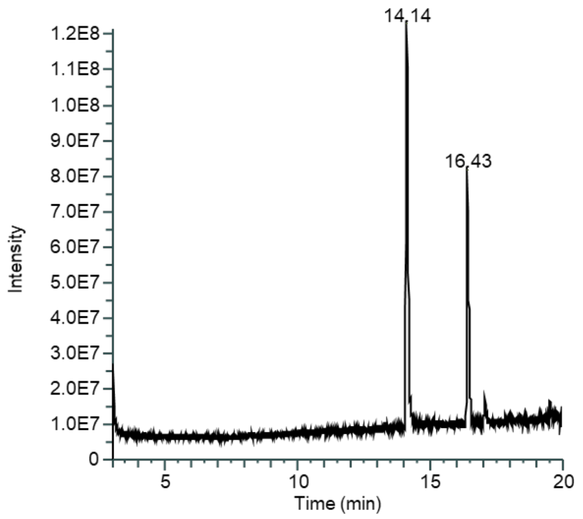


ms

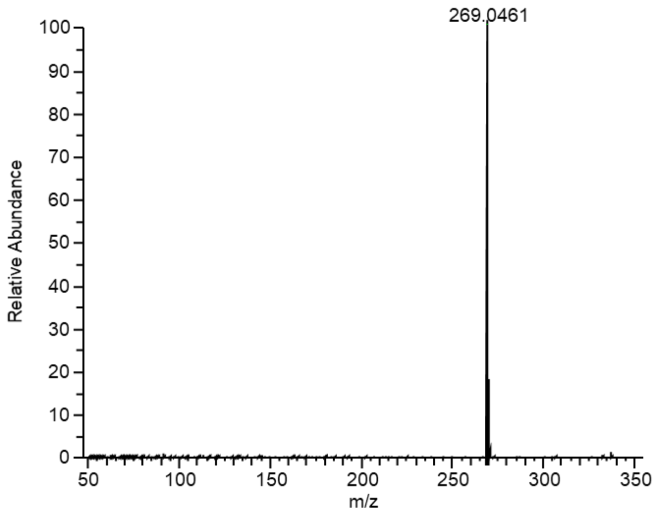


ms2

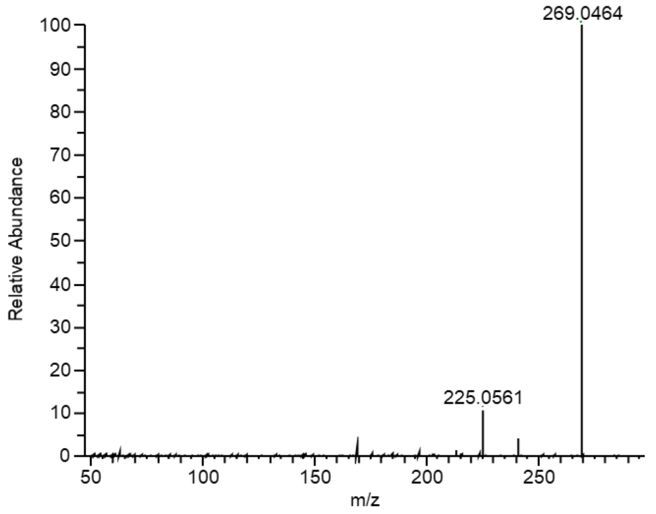


Fig. S5

Aoki et al.

TIC

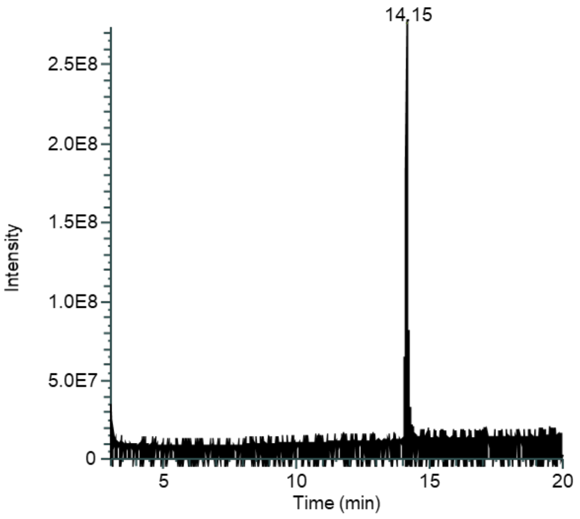


ms

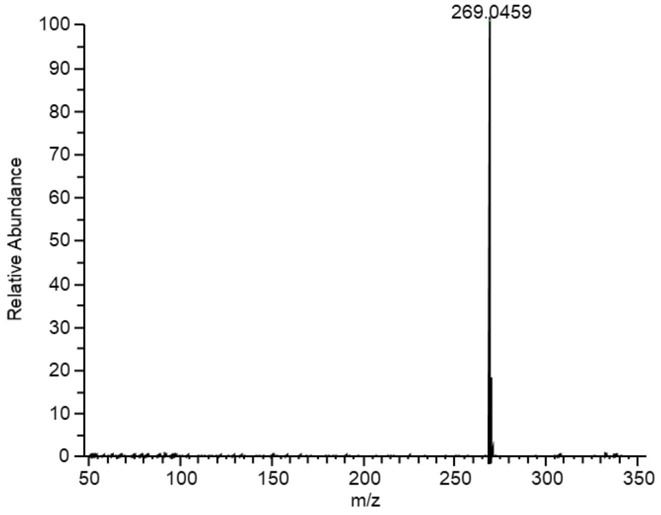


ms2

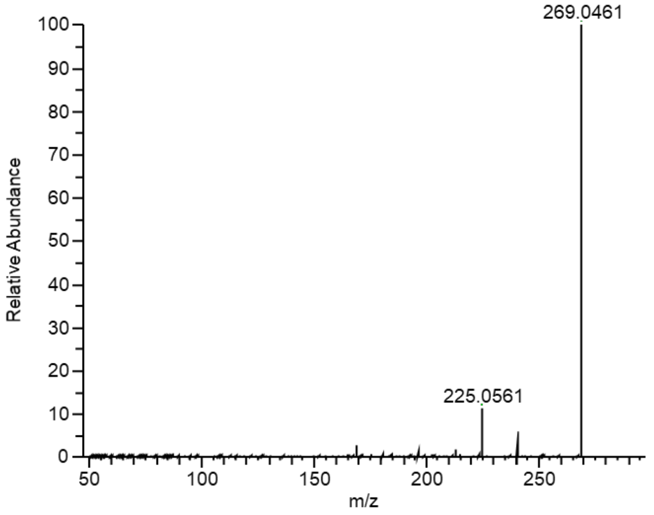


Fig. S6

Aoki et al.

TIC

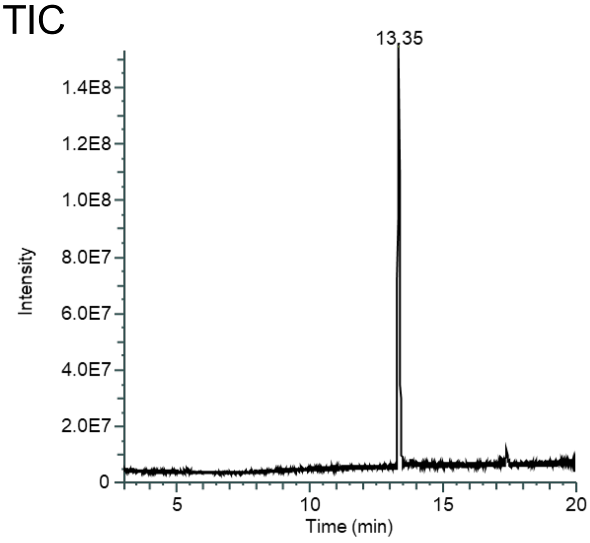


ms

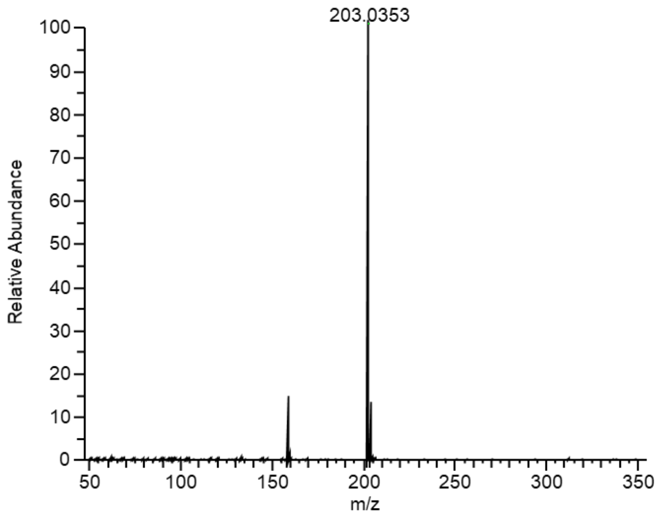


ms2

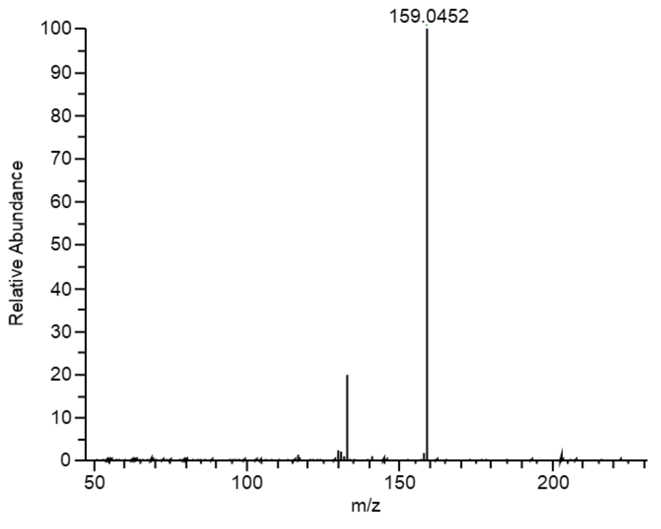


Fig. S7

Aoki et al.

(a)

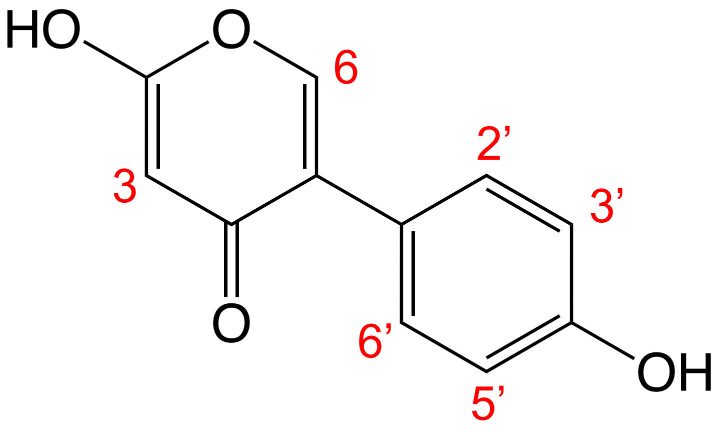


2-hydroxy-5-(4-hydroxyphenyl)-4*H*-pyran-4-one

(b)

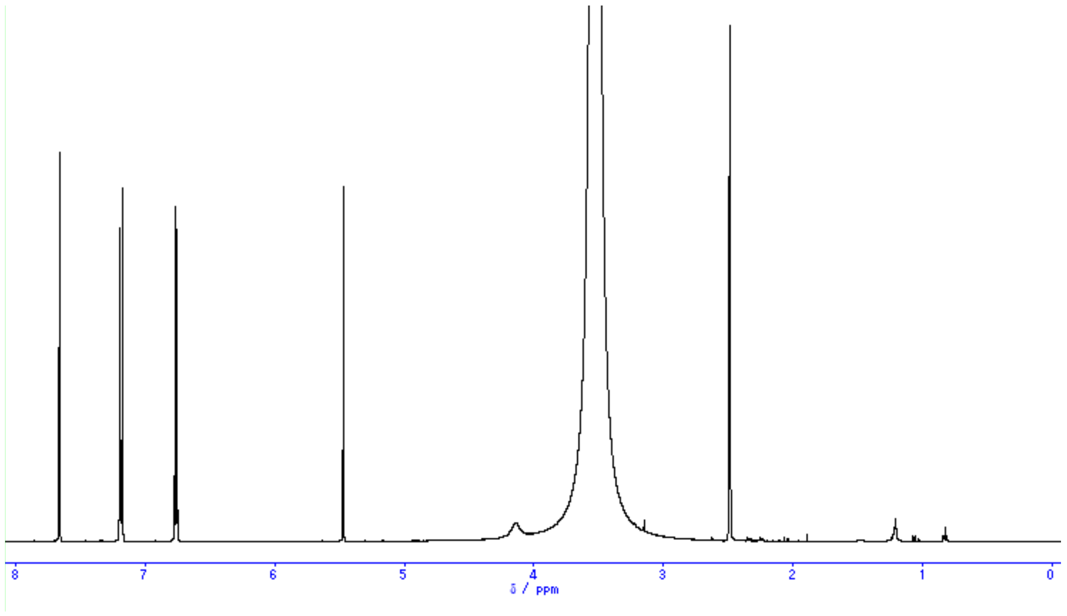


Fig. S8

Aoki et al.

**(a)** IFCA product

ms

ms2

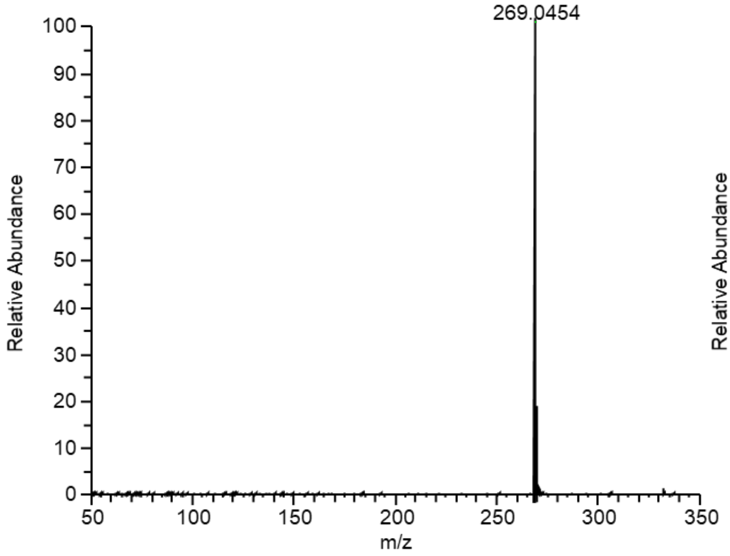


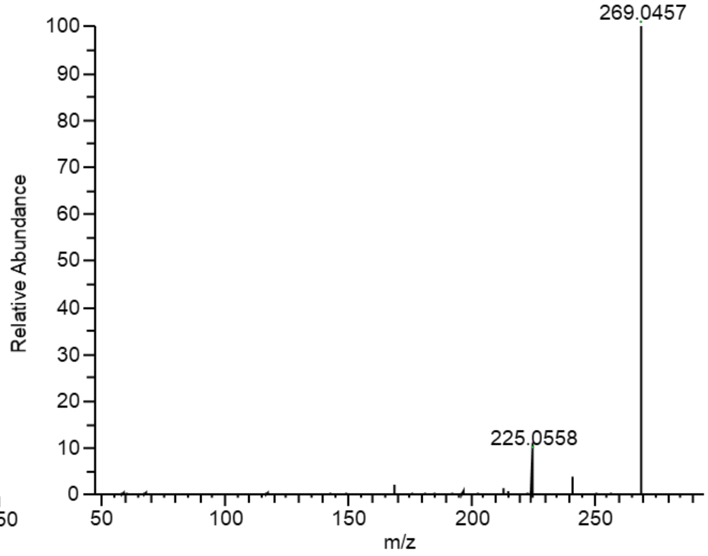


**(b)** 8-Hydroxydaidzein standard

ms

ms2

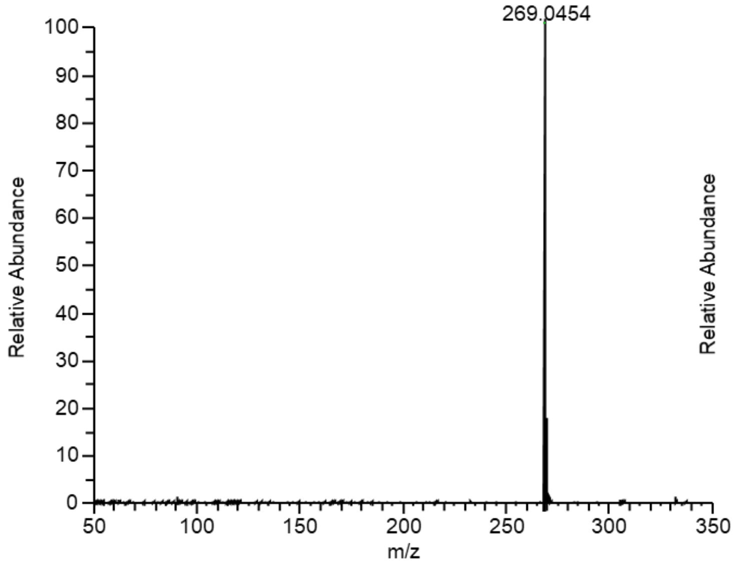


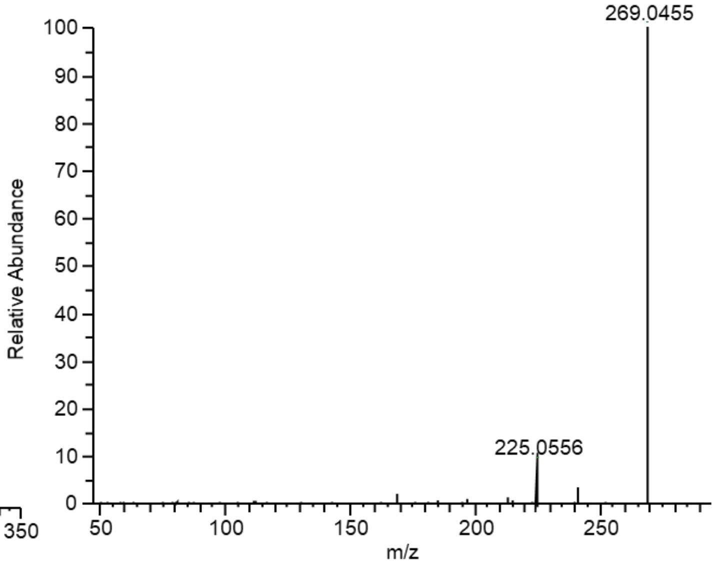


Fig. S9

Aoki et al.

IFCA + daidzein

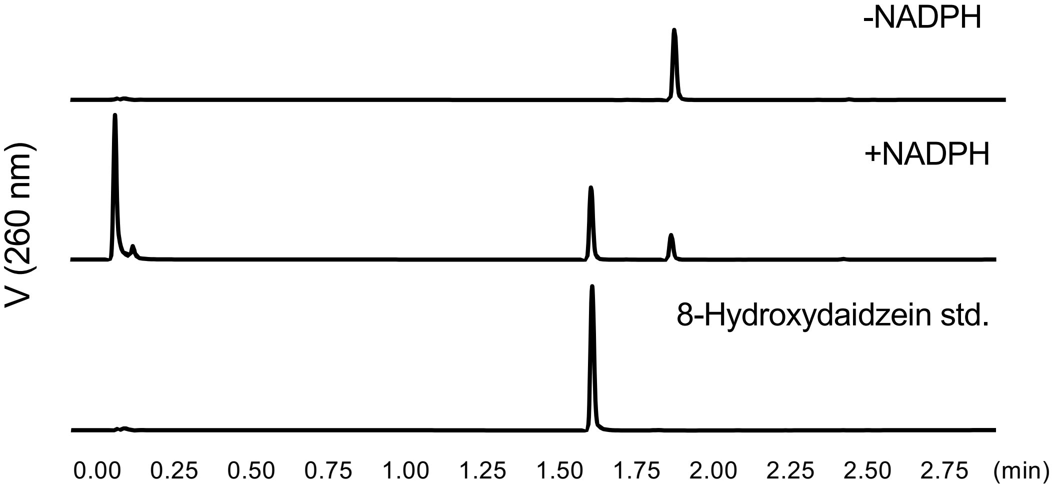


Fig. S10

Aoki et al.

### Reactive substrates

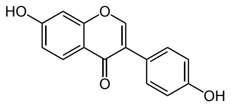


Daidzein

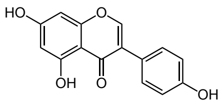


Genistein

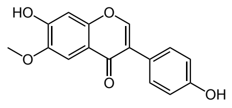


Glycitein

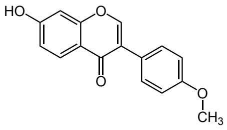


Formononetin

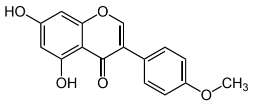


Biochanin A

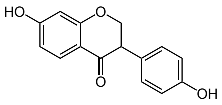


Dihydrodaidzein

---

### Un-reactive substrates

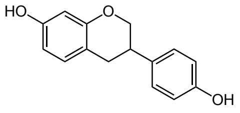


Equol

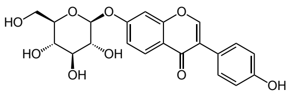


Daidzin

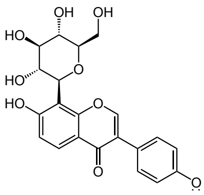


Puerarin

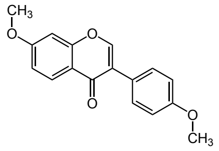


4',7-Dimethoxyisoflavone

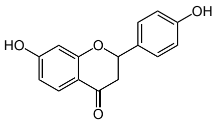


Liquiritigenin

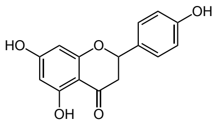


Naringenin

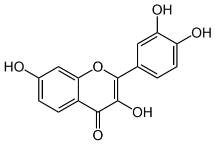


Quercetin

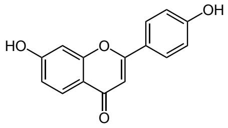


4′,7-
Dihydroxyflavone

Fig. S11

Aoki et al.

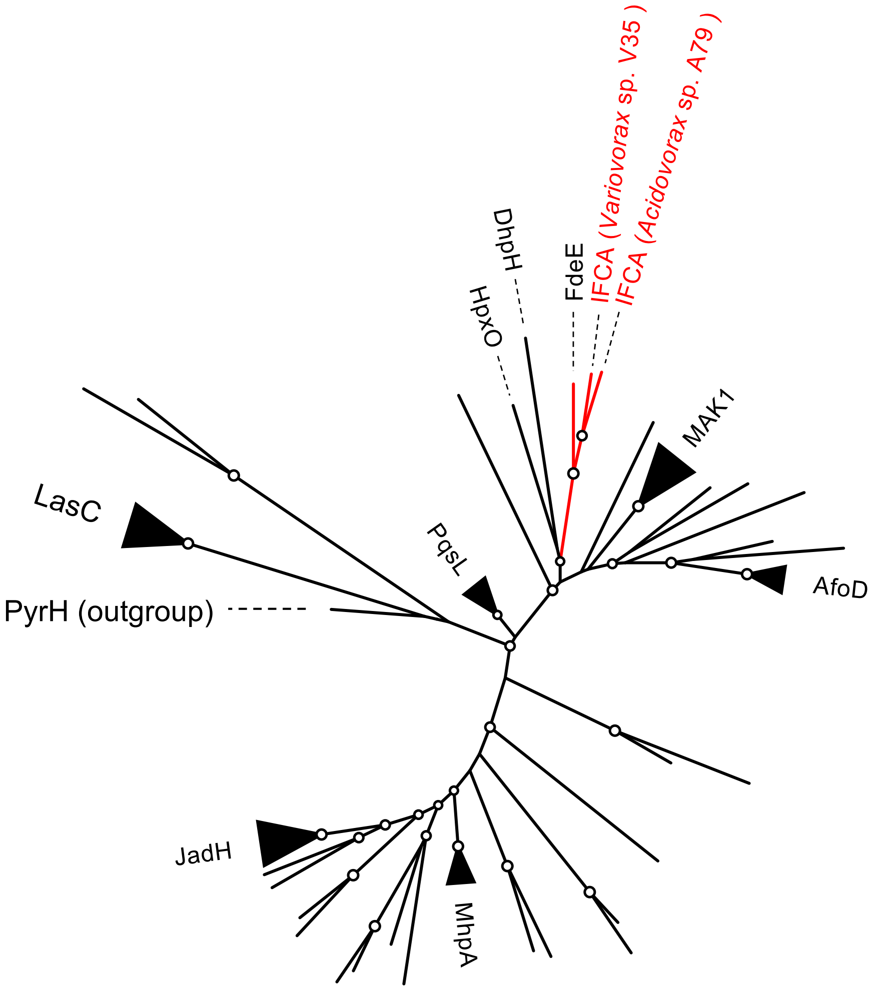


Fig. S12

Aoki et al.

**(a)** IFCB product

ms

ms2

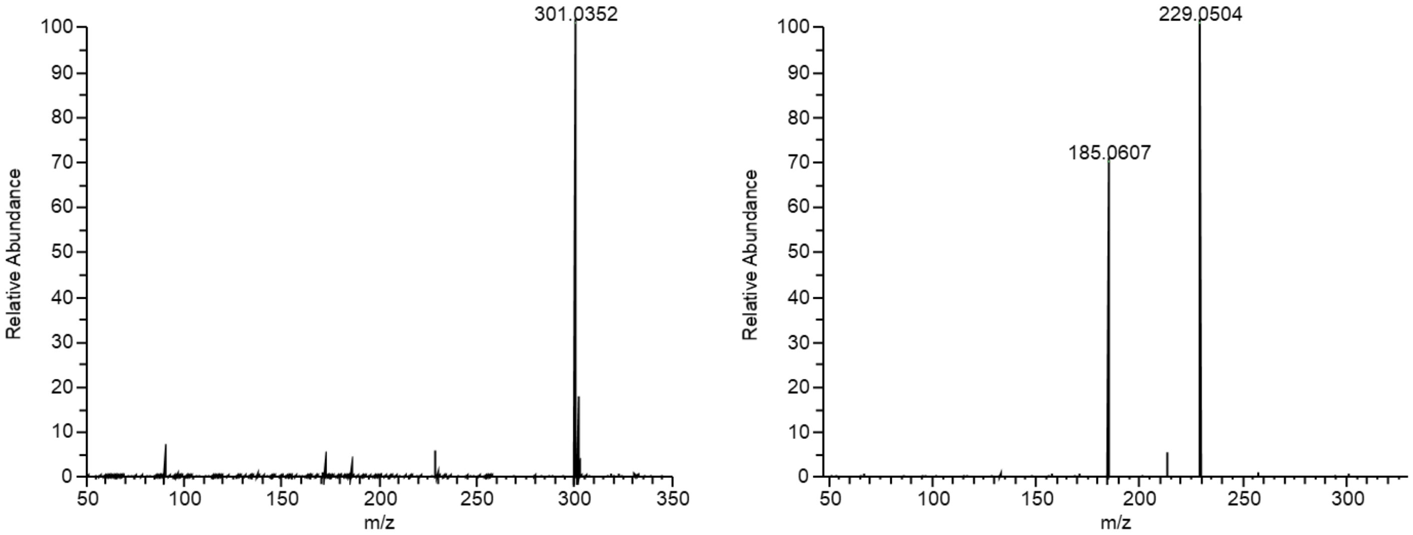


**(b)** Compound $X_2$

ms

ms2

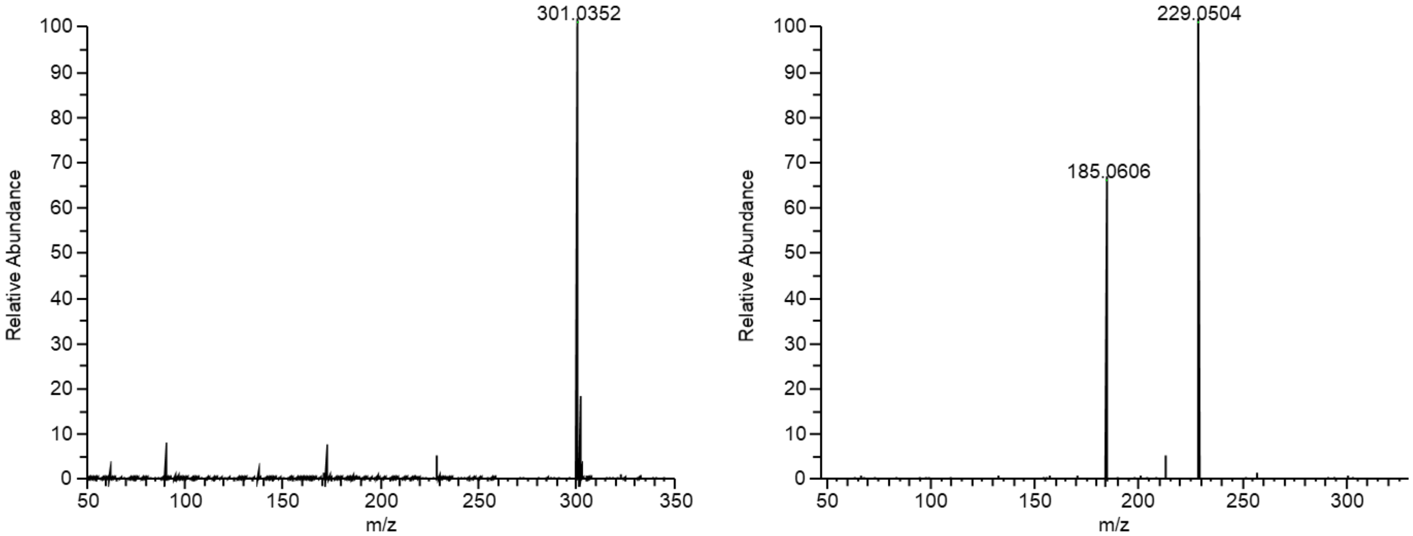


Fig. S13

Aoki et al.

#### (a) IFCD1 product 1

ms

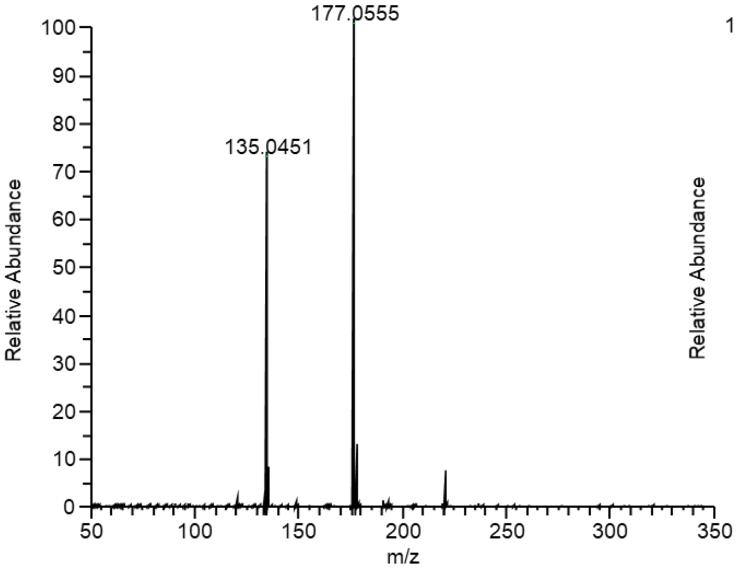


ms2

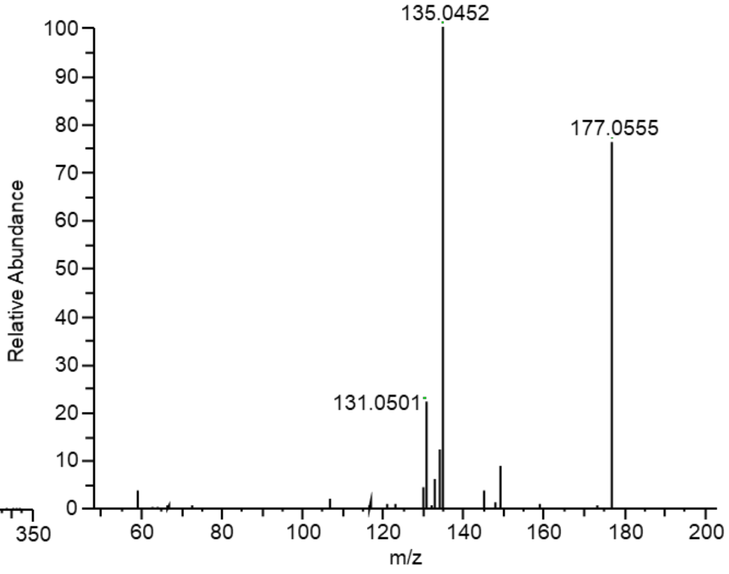


#### (b) IFCD1 product 2

ms

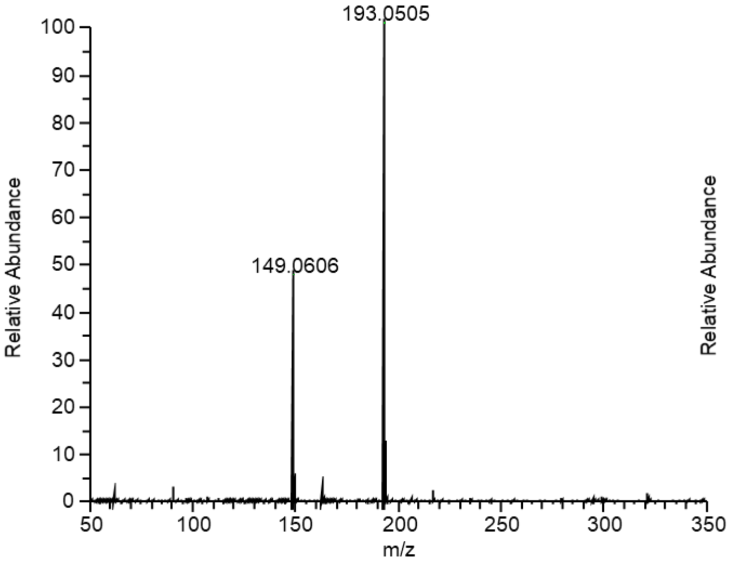


ms2

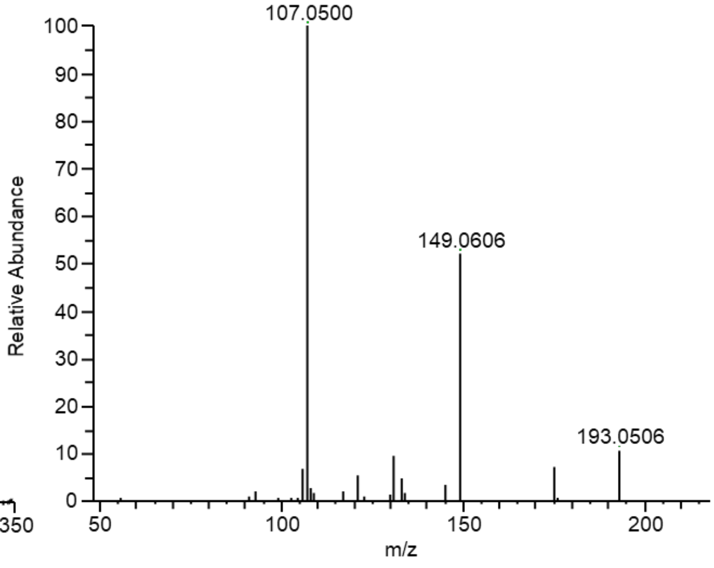


Fig. S14

Aoki et al.

**(a)** IFCD2 product 1

ms

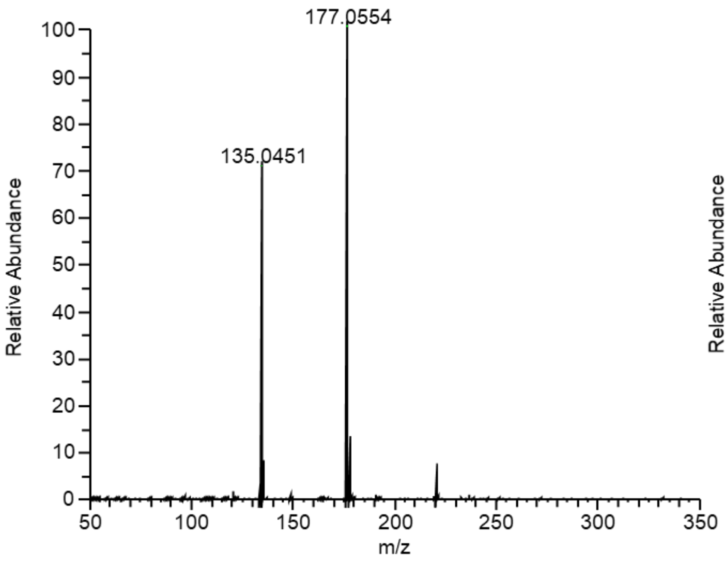


ms2

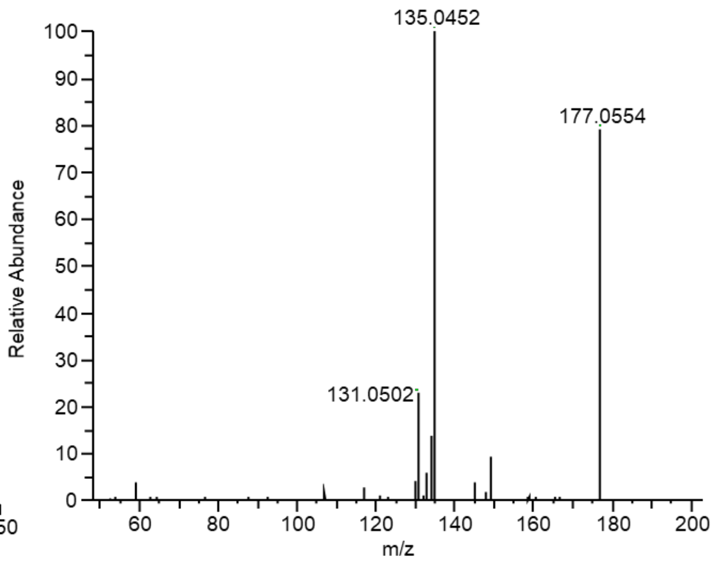


**(b)** IFCD2 product 2

ms

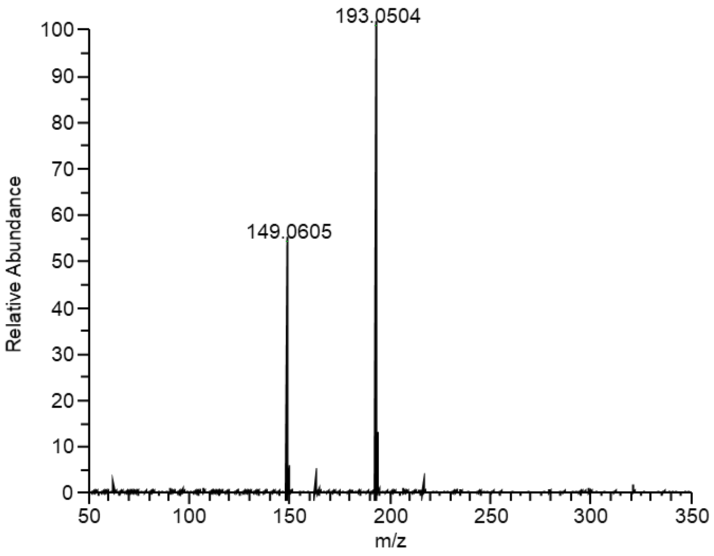


ms2

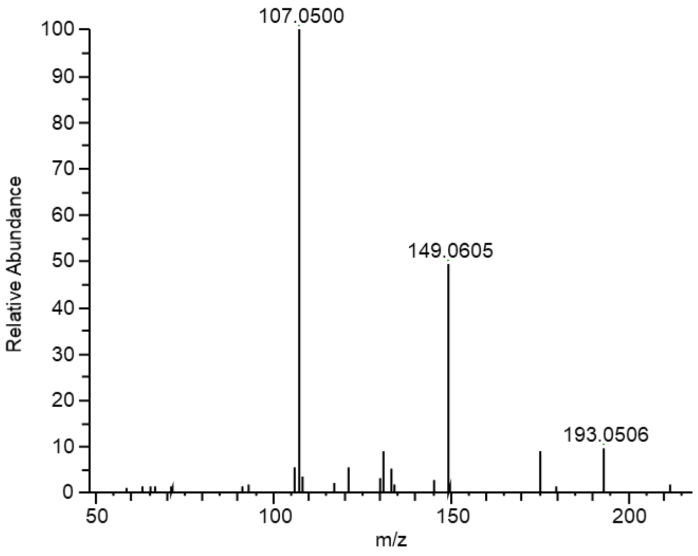


Fig. S15

Aoki et al.

**(a)** Compound $X_3$-$d_2$

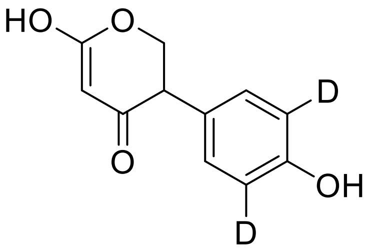


**(b)** IFCD1 product 1

ms

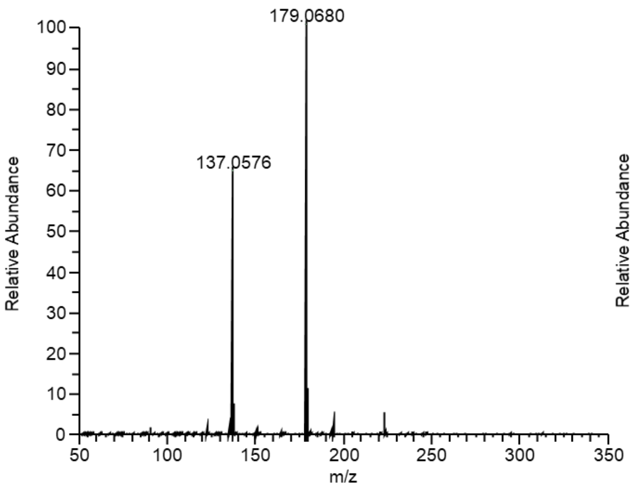


ms2

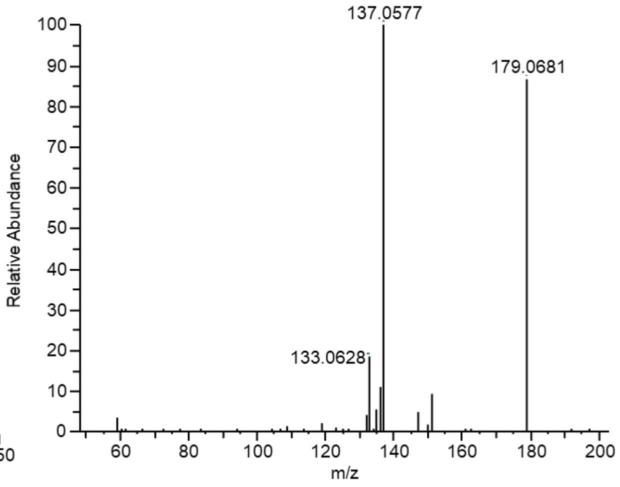


**(C)** IFCD1 product 2

ms

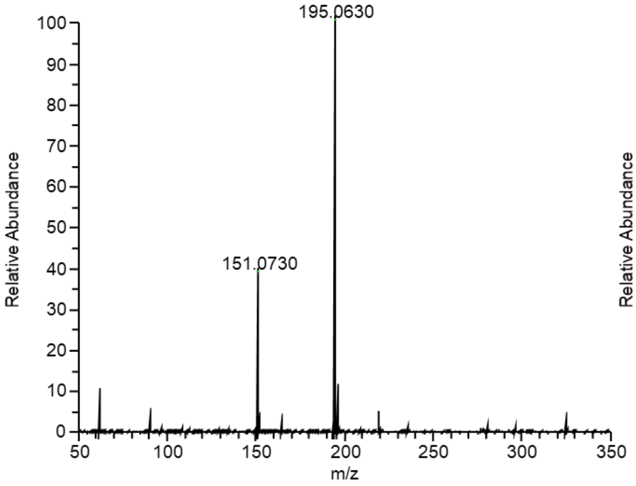


ms2

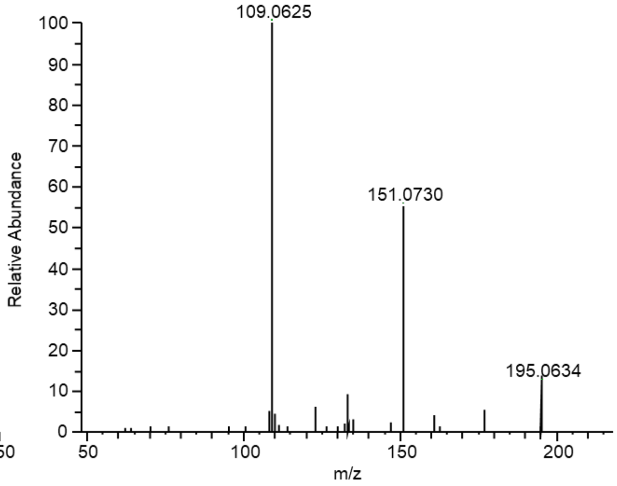


Fig. S16

Aoki et al.

**(a)** Compound $X_3$-$d_4$

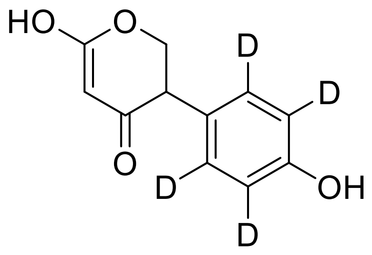


**(b)** IFCD1 product 1

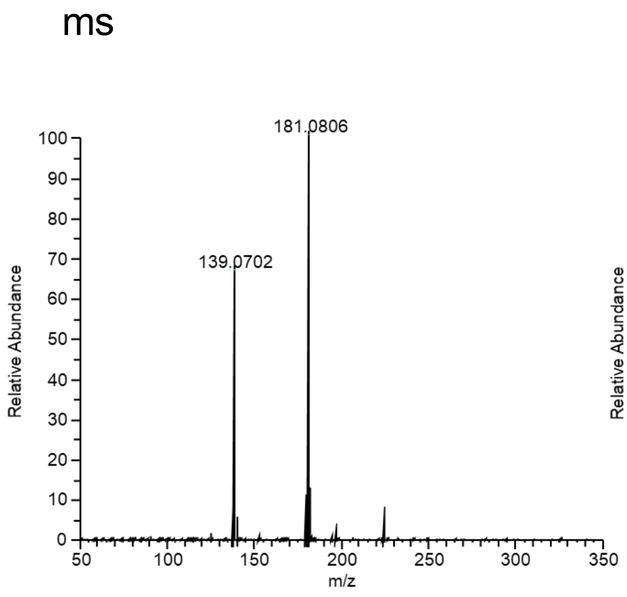


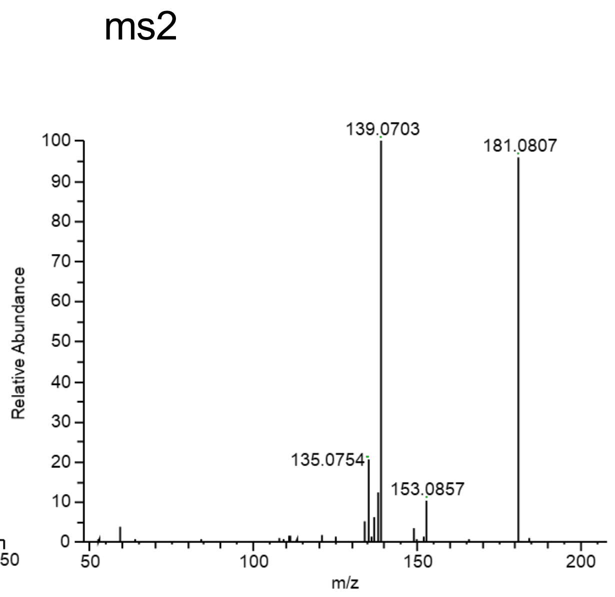


**(C)** IFCD1 product 2

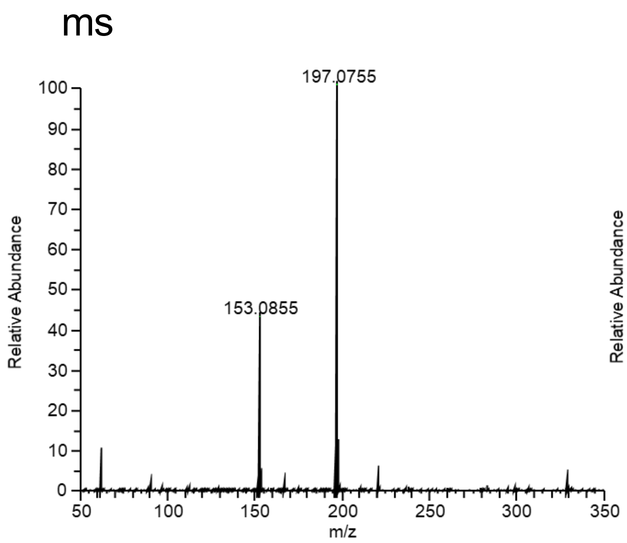


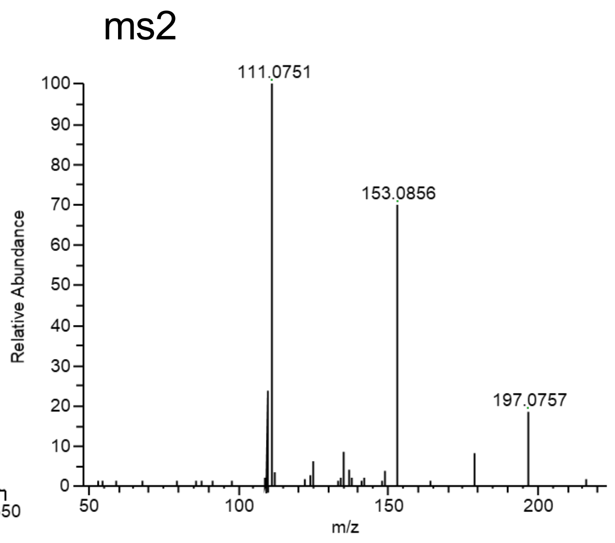


Fig. S17

Aoki *et al.*,

**(a)** $C_{10}H_{10}O_3$

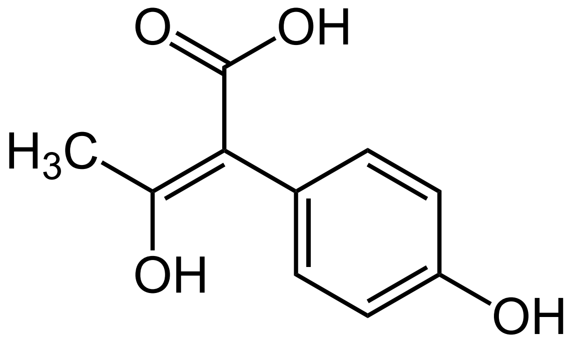

**(b)** $C_{10}H_{10}O_4$

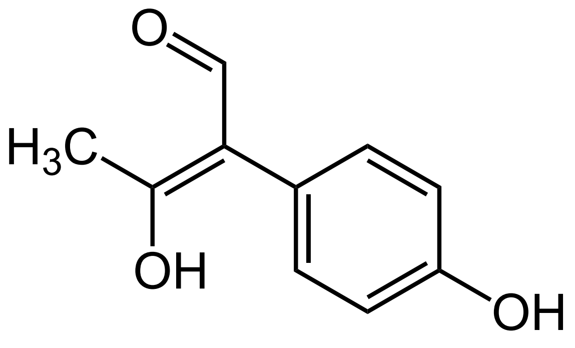

Fig. S18

Aoki et al.

### Meta-cleavage of catechol pathway (M00569)

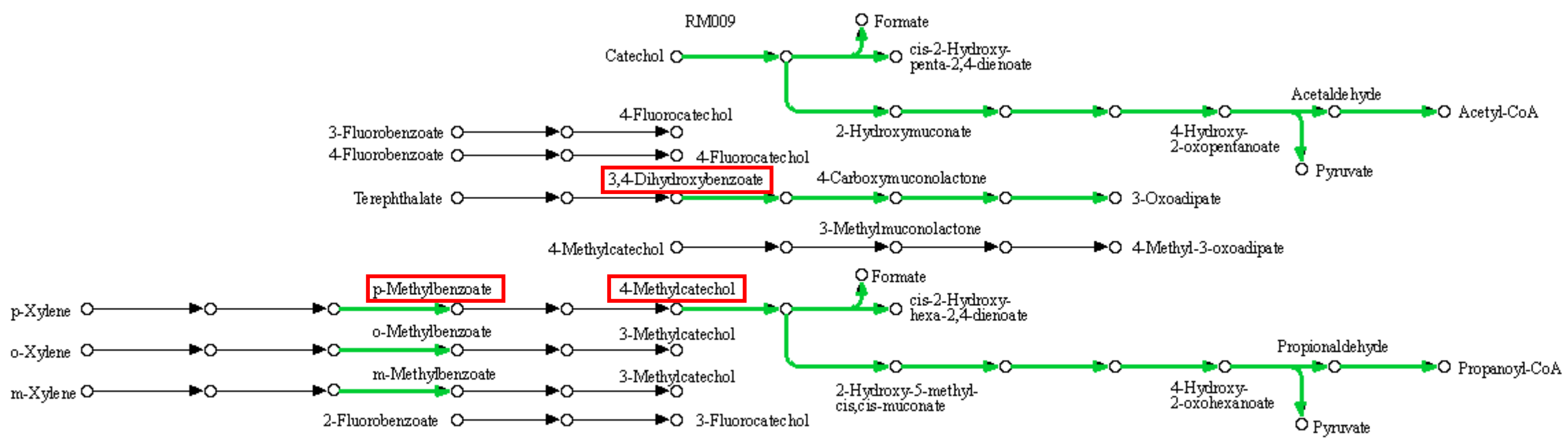


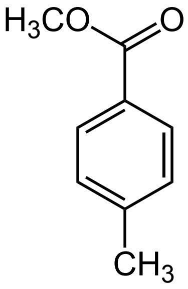


p-Methylbenzoate

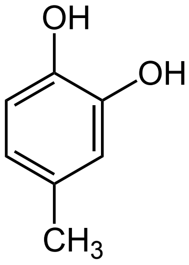


4-Methylcatechol

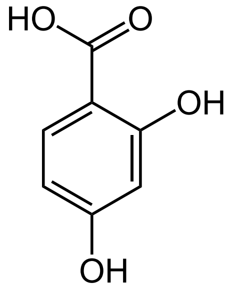


3,4-Dihydroxybenzoate

Aoki et al.

Fig. S19

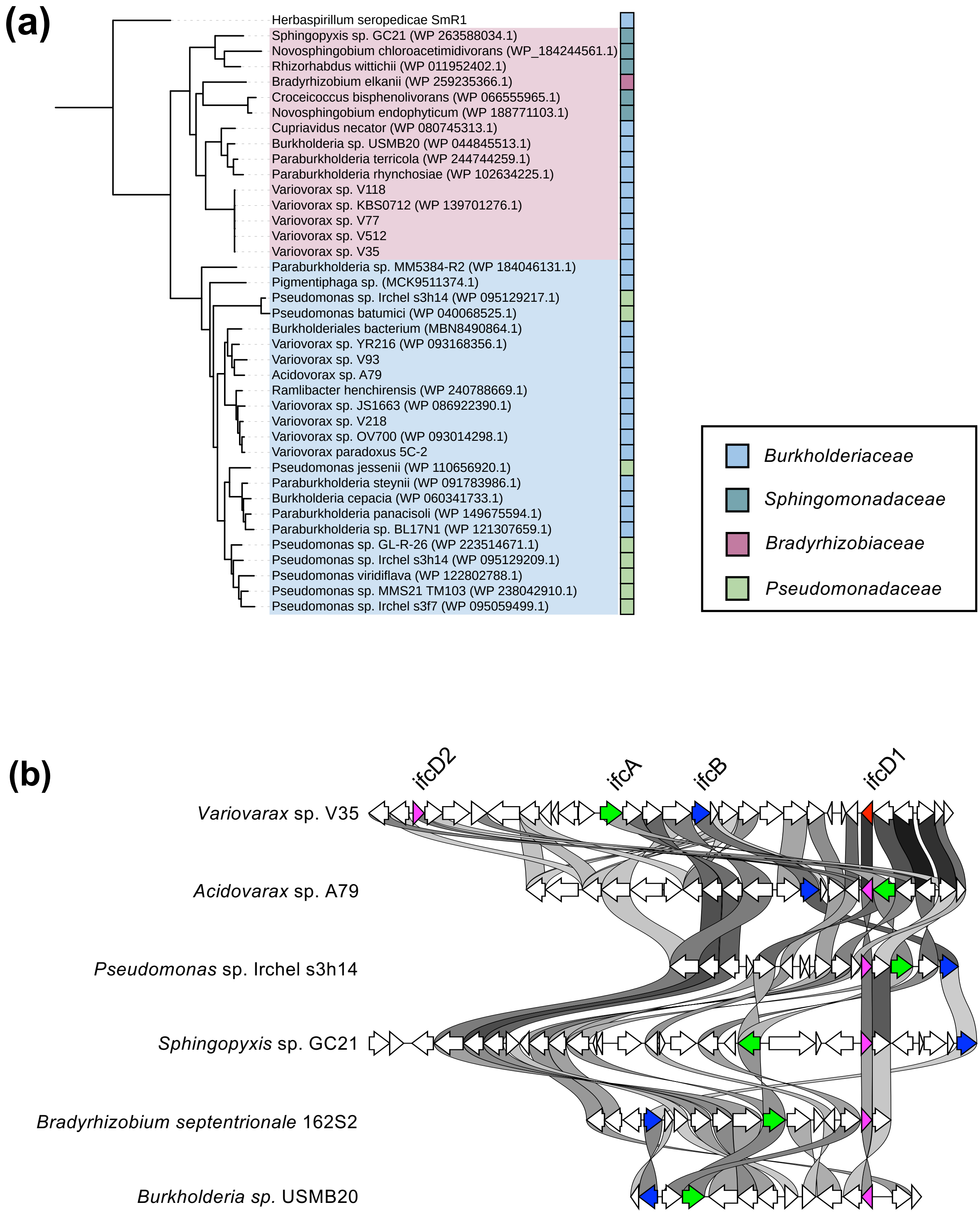


Fig. S20

Aoki et al.
